# FDA-approved drug library screen identifies antidepressants, antimicrobials, anti-COPD, and anti-CVD agents as blockers of NLRP3 inflammasome and sepsis in a sex-dependent manner

**DOI:** 10.64898/2026.03.05.709979

**Authors:** Kara Timinski, Ashutosh Prince, Kalash Neupane, Swati Sharma, Nilam Bhandari, Mariam R Khan, Yavar Shiravand, C. Alicia Traughber, Zackery Raquepaw, Kailash Gulshan

## Abstract

The NLRP3 inflammasome pathway is central to host defense, but dysregulated activation of inflammasomes promotes diseases associated with metabolic syndrome (diabetes, obesity, CVD, MASLD), neurodegenerative diseases (Alzheimer’s, Parkinson’s), autoinflammatory conditions (CAPS, gout), and respiratory illnesses (asthma/COPD, COVID-19). Therapeutic modulation of NLRP3 is challenging as it requires selective blockade of detrimental inflammasome activation without broadly suppressing innate immunity. Here, we used a phenotypic screen in THP1-ASC-GFP monocytes to identify FDA-approved drugs that can block LPS-induced priming of the NLRP3 inflammasome or inhibit NLRP3 assembly without disrupting upstream priming. Various classes of drugs, such as antidepressants (Fluoxetine, Duloxetine), antihypertensives (Irbesartan, Amlodipine, Nebivolol), antidiabetics (Rosiglitazone), β-adrenergic agonists (Salmeterol), antimalarials (Mefloquine), antifungals (Azoles, Ciclopirox), and antivirals (Saquinavir, Remdesivir), were identified as potent blockers of either priming or assembly of the NLRP3 inflammasome. Secondary validation demonstrated that several compounds suppressed NF-κB activation, reduced LPS binding to immune cells, decreased pro-inflammatory cytokine production, enhanced efferocytic capacity of macrophages, and enhanced autophagy *in vitro* and *in vivo*. Mechanistic analyses further revealed drug-specific effects on lysosomal biogenesis, mitochondrial morphology, and mitochondrial reactive oxygen species. In murine models of acute inflammation and endotoxemia, selected compounds attenuated NLRP3 inflammasome activation, reduced systemic and local inflammatory cytokines, limited tissue injury, and improved survival following LPS-induced sepsis, with efficacy varying by sex. Further studies in primary human cells and *in vivo* disease models are needed to assess the repurposing and therapeutic relevance of identified drugs.

## Introduction

The NLRP3 inflammasome is a central regulator of innate immune responses and contributes to the pathogenesis of sepsis, atherosclerosis, metabolic disorders, neurodegeneration, and numerous chronic inflammatory diseases^1–3^. Under homeostatic conditions, NLRP3 is degraded via polyubiquitination^4,5^. Upon receiving a “priming” signal (Signal 1), a transcriptional cascade is initiated via nuclear localization of NF-kB, leading to upregulation of several genes, such as NLRP3 and pro-IL-1β/IL-18. A secondary “activation” signal (Signal 2), such as potassium efflux, extracellular ATP, or lysosomal damage, triggers NLRP3 inflammasome assembly and cleavage of procaspase-1. Activated caspase-1 executes two critical proteolytic functions: cleaving the inactive precursors pro-IL-1β and pro-IL-18 into their mature forms, and cleaving Gasdermin D (GSDMD). GSDMD was identified as the ultimate executioner of pyroptotic cell death^6–8^. While essential for host defense, aberrant or chronic NLRP3 overactivation drives numerous pathologies, including cardiovascular diseases (CVD), chronic obstructive pulmonary disease (COPD), metabolic syndromes, neurodegenerative disorders, and cancer^3^. Recent studies also highlighted the role of inflammasome-mediated cytokine storm in COVID-19 and sepsis^9–11^. The initial host response to microbial infection triggers a robust pro-inflammatory response necessary for pathogen clearance. However, this response becomes dysregulated, leading to systemic inflammation and organ damage. The interconnectedness of TLR and NLRP3 pathways makes targeting both an attractive therapeutic strategy for mitigating the uncontrolled inflammation characteristics of sepsis. This intricate relation between priming (TLR2/4) and activation of the NLRP3 inflammasome highlights the need for identifying drugs that can specifically target these steps. Existing therapies, such as the biologics Anakinra and Canakinumab, block downstream IL-1 activity but fail to prevent the full spectrum of inflammasome-mediated damage, including pyroptosis and the release of other inflammatory mediators. Previous studies showed promising results with MCC950, a highly selective small-molecule inhibitor of the NLRP3 inflammasome. MCC950 has demonstrated remarkable efficacy in several animal disease models, ranging from atherosclerosis, Alzheimer’s, IBD, and rescuing neonatal lethality in models of cryopyrin-associated periodic syndrome (CAPS)^12^. While MCC950 showed promise in preclinical trials, its clinical development was reportedly halted in Phase II trials for rheumatoid arthritis due to concerns over liver toxicity. Although several NLRP3-targeting drugs are or have been in clinical trials^13^, drug repurposing offers an attractive alternative because FDA-approved compounds possess well-characterized pharmacokinetic and safety profiles that can accelerate clinical translation. In the present study, we identified multiple clinically approved drugs capable of inhibiting distinct stages of inflammasome activation and demonstrated that these compounds not only suppress inflammatory signaling but also activate endogenous pathways involved in inflammation resolution and cellular homeostasis.

Several established clinical agents—including antidepressants, antimalarials, antifungals, antivirals, and medications for COPD and CVD—exhibit robust inhibitory activity against NLRP3 priming/activation. We classified the candidates into two categories: 1) drugs blocking LPS priming and 2) drugs blocking NLRP3 inflammasome assembly (ASC speck formation). The drugs that block LPS priming or ASC speck formation without affecting the expression of inflammasome components (NLRP3, ASC, procaspase-1) were further tested for mechanistic insights and efficacy in blocking *in vivo* inflammasome assembly and LPS-induced mortality in mice. These data provide a strong rationale for repurposing these diverse clinical agents as an adjuvant therapeutic along with standard care to treat a variety of NLRP3-associated inflammatory conditions.

## Results

### Screening of FDA-approved drugs identifies several classes of drugs blocking LPS priming

To identify clinically approved drugs capable of inhibiting the priming or assembly of the NLRP3 inflammasome, a microscopy-based screen using an FDA-approved drug library and THP1-ASC-GFP monocytic cells (***Fig. 1A***) was employed. The THP1-ASC-GFP cells stably express a gene encoding an ASC-GFP fusion protein under the control of an NF-κB-inducible promoter. Control cells show a basal and weak GFP signal, while LPS treatment robustly induced ASC-GFP expression, as evidenced by increased GFP signal in cells (***Fig. 1B***). Addition of Nigericin to LPS-treated cells leads to robust NLRP3 inflammasome assembly, with a single ASC-GFP puncta formed in most of the cells (***Fig. 1B***). Primary screening revealed a subset of compounds that significantly attenuated LPS-induced priming verses control cells (***Fig. 1C***), including Febuxostat (***Fig. 1D***), Dasatinib (***Fig. 1E***), Ciclopirox (***Fig. 1F***), Mefloquine (***Fig. 1G***), Fluoxetine (***Fig. 1H***), ABT (***Fig. 1I***), Amlodipine (***Fig. 1J***), Bepridil (***Fig. 1K***), Nebivolol (***Fig. 1L***), Tamoxifen (***Fig. 1M***), Remdesivir (***Fig. 1N***), Aripiprazole (***Fig. 1O***), Duloxetine (***Fig. 1P***), Bazedoxifene (***Fig. S1***), and FTY720 (***Fig. S1***). All identified drugs showed >50% reduction in ASC-GFP fluorescence (***Fig. 1Q***).

**Fig. 1:**
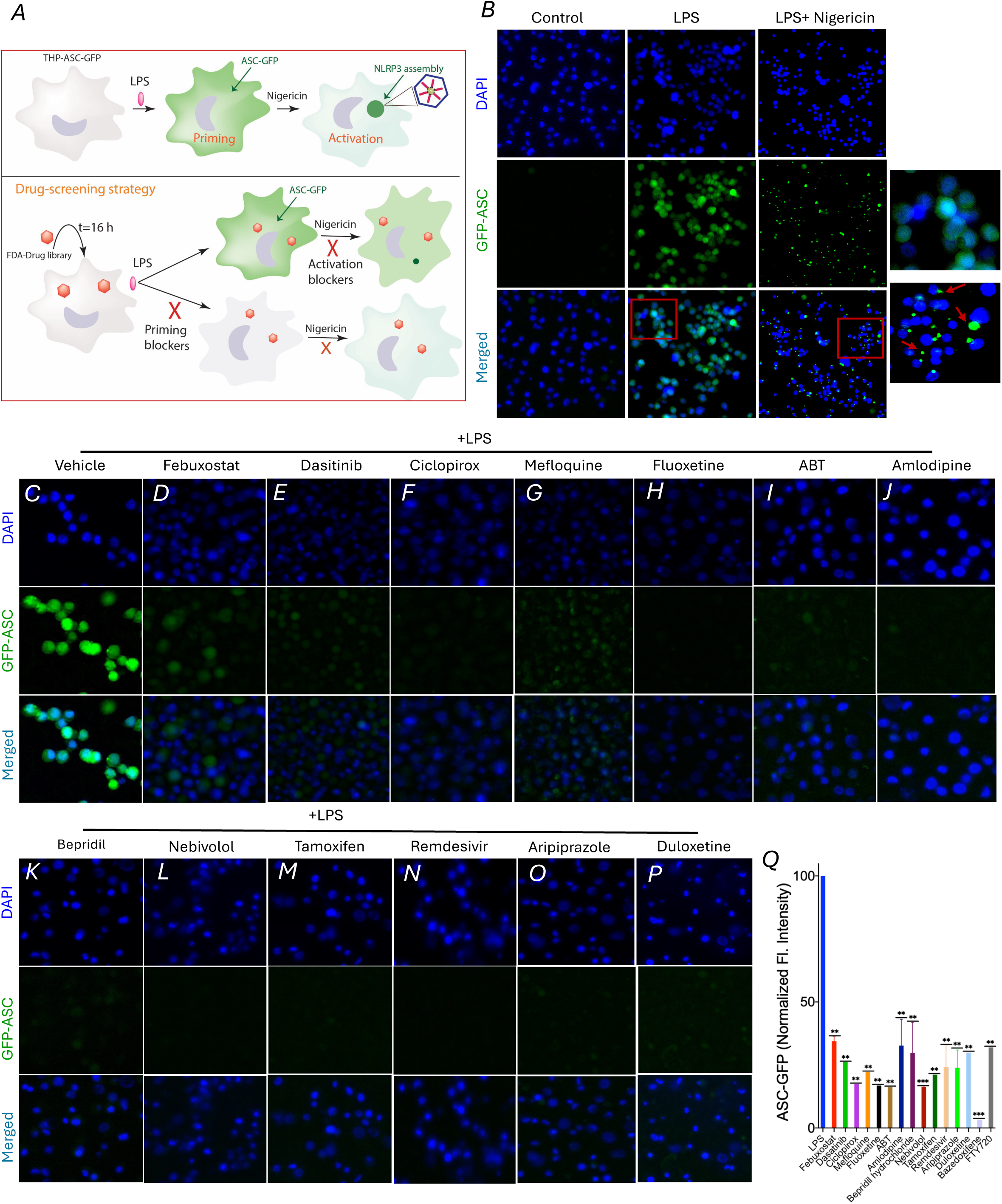
Screening of FDA-approved drugs identifies several classes of drugs blocking LPS priming. (***A***) Schematic diagram showing the FDA-approved drug library screen with the human monocytic THP-1ASC-GFP cell line. Cells were pretreated with vehicle or 20μM of the indicated drug for 16h, followed by LPS priming for 2h and 10μM Nigericin treatment for 1h to induce NLRP3 inflammasome assembly (ASC puncta formation). (***B***) Control THP-1-ASC-GFP cell line stained with DAPI showing low-level basal expression of ASC-GFP. Cells treated with LPS showing induced ASC-GFP expression (central row), and cells treated with LPS + Nigericin showing ASC-GFP puncta formation (right row). (***C***) Control cells, or cells treated with Febuxostat (***D***), Dasatinib (***E***), Ciclopirox (***F***), Mefloquine (***G***), Fluoxetine (***H***), ABT (***I***), Amlodipine (***J***), Bepridil (***K***), Nebivolol (***L***), Tamoxifen (***M***), Remdesivir (***N***), Aripiprazole (***O***), or Duloxetine (***P***), were primed with LPS for 4h (instead of 2h for more robust ASC-GFP expression) and subjected to fluorescent microscopy. (***Q***) Quantification of GFP fluorescence signal in cells treated with LPS ± various drug pretreatments for 16 h (N=3 panel, with each panel ∼100 cells, values are mean ± SD, significance was determined by t-test, comparing each column with + LPS sample, with, ** indicating p<0.01, and *** indicating p<0.001).

### Effect of inflammasome priming blockers on downstream signaling pathways

Consistent with inhibition of priming, pretreatment of RAW-ASC or THP-1 macrophages with validated compounds such as Febuxostat, Ciclopirox, Mefloquine, and Fluoxetine significantly reduced NF-kB nuclear localization verses control LPS-treated cells (***Fig. 2A-D***). In RAW-ASC cells, most of the priming blockers showed more than 50% reduction in LPS-induced NF-kB nuclear localization (***Fig. 2B***, n ≥ 50 for all, mean ± SD, * indicates p <0.05, ** indicates p<0.01, *** indicates p<0.001 by t-test). Similar results were found in THP-1 macrophages (***Fig. 2C, Fig. 2D***), with the only exception of Febuxostat, which showed a trend toward reduced NF-kB nuclear localization but did not reach statistical significance (***Fig. 2C, 2D***). Furthermore, priming blocking drugs, such as Ciclopirox and Mefloquine, markedly reduced LPS-induced upregulation of pro-inflammatory mediators associated with inflammasome assembly, including NLRP3 and pro-IL-1β (***Fig. 2E-2H)***. Importantly, these compounds did not affect basal ASC-GFP expression in unstimulated or LPS-induced cells, supporting a selective effect on LPS-induced inflammatory signaling (***Fig. 2E***). Similarly, Febuxostat treatment led to a significant reduction in LPS-induced expression of NLRP3 and pro-IL-1β (***Fig. 2I, 2J***).

**Fig. 2:**
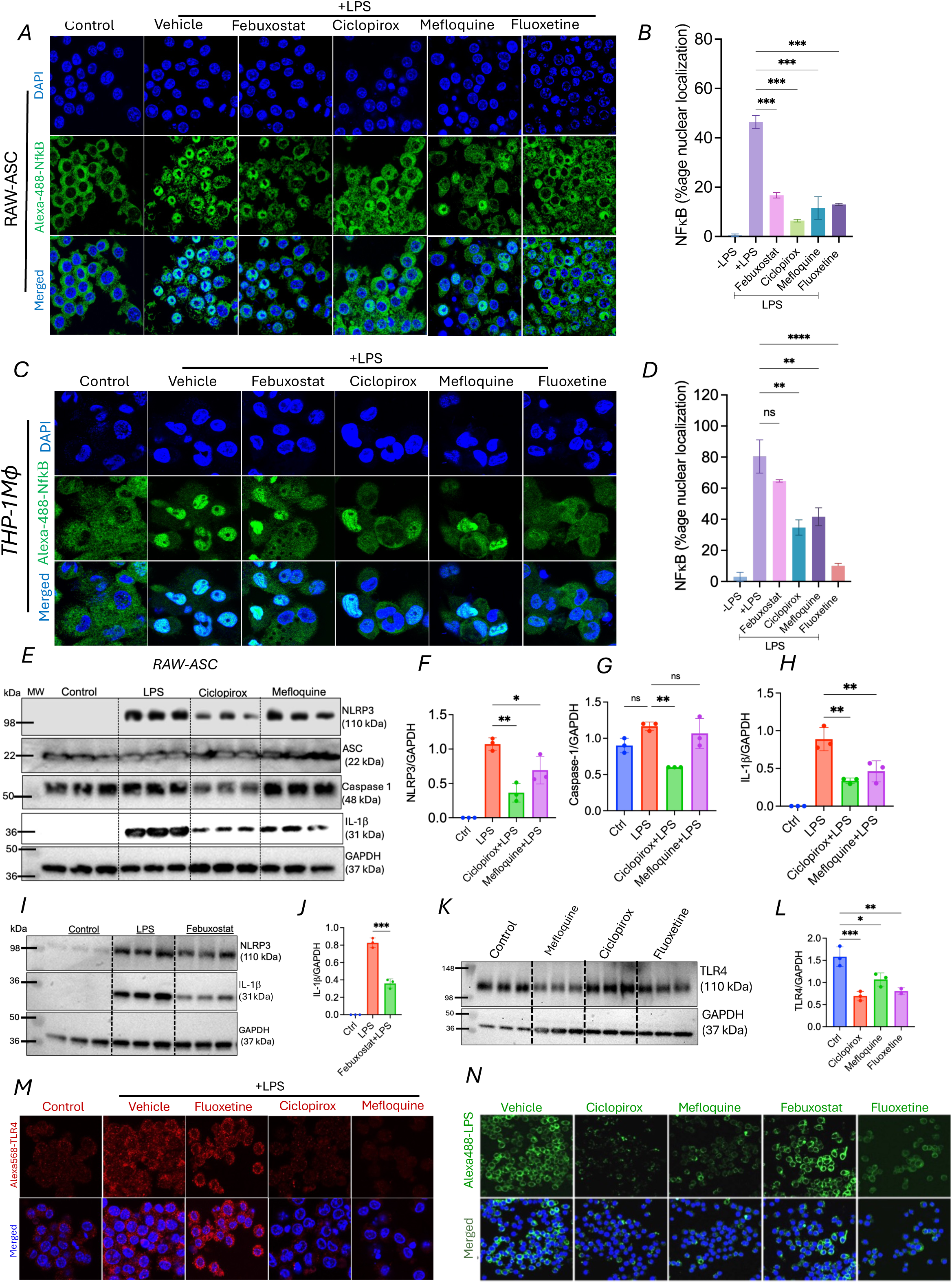
Effect of inflammasome priming blockers on downstream signaling pathways. (***A***) RAW-ASC cells were treated with vehicle or 20μM of the indicated drug for 16h, followed by 1μg/ml LPS priming for 2h. Cells were washed, fixed, and subjected to indirect immunofluorescence using rabbit anti-NF-kB (p65) antibody and goat anti-rabbit Alexa-488 antibody. DAPI was used to stain nucleus. (***B***) Graph bars showing quantification of % of cells showing nuclear NF-kB localization. Values displayed as mean ± SD, significance was determined by t-test, comparing each column with + LPS sample, with, *** indicating p<0.001. (***C***) THP-1 cells were differentiated into macrophages by PMA treatment and treated with vehicle or 20μM of the indicated drug for 16 h, followed by LPS priming. Cells were washed and fixed and subjected to indirect immunofluorescence using rabbit anti-NF-kB (p65) antibody and goat anti-rabbit Alexa-488 antibody. DAPI was used to stain nucleus. (***D***) Graph bars showing quantification of % of cells showing nuclear NF-kB localization. Values displayed as mean ± SD, significance was determined by t-test, comparing each column with + LPS sample, with, * indicating p<0.05, ** indicating p<0.01, *** indicating p<0.001, **** indicating p<0.0001. (***E***) RAW-ASC cells were treated with vehicle or 20μM of Ciclopirox or Mefloquine for 16h, followed by LPS priming for 4h. Western blots were performed using antibodies against NLRP3, ASC, Caspase1, or IL-1β, with GAPDH used as loading control. Graph bars showing quantification using GAPDH ratio to (***F***) NLRP3, (***G***) Procaspase-1, and (***H***) IL-1β. Values displayed as mean ± SD, significance was determined by t-test, comparing each column with + LPS sample, with, * indicating p<0.05, ** indicating p<0.01, ns indicates non-significant. (***I***) RAW-ASC cells were treated with vehicle or 20μM of Febuxostat for 16h, followed by LPS priming for 4h. Western blots were performed using antibodies against NLRP3 or IL-1β, with GAPDH used as loading control. (***J)*** Graph bars showing quantification using GAPDH ratio to IL-1β. Values displayed as mean ± SD, significance was determined by t-test, comparing each column with + LPS sample, with, *** indicating p<0.001. (***K***) RAW-ASC cells were treated with vehicle or 20μM indicated drugs for 16 h, and western blots were performed using antibodies against TLR4, with GAPDH used as loading control. (***L)*** Graph bars showing quantification using GAPDH ratio to TLR4. Values displayed as mean ± SD, significance was determined by t-test, with * indicating p<0.05, ** indicating p<0.01, *** indicating p<0.001. (***M***) RAW-ASC cells were treated with vehicle or 20μM indicated drugs for 16 h, followed by LPS priming for 4h. Cells were washed and fixed, and subjected to indirect immunofluorescence using rabbit anti-TLR4 antibody and goat anti-rabbit Alexa-568 antibody. DAPI was used to stain nucleus. (***N***) RAW264.7 cells were treated with vehicle or 20μM indicated drugs for 16h, followed by incubation with (10ug/mL) Alexa-488 labeled LPS for 2 hour, followed by washing and visualization under a fluorescent microscope. DAPI was used to stain nucleus.

As TLR4 plays a major role in relaying LPS signaling, we tested if any of the selected drugs affected basal TLR4 levels. As shown in ***Fig. 2K, 2L***, the expression levels of TLR4 were significantly reduced in cells treated with Mefloquine, Ciclopirox, and Fluoxetine. Next, we determined if selected drugs also affect LPS-induced enhancement of TLR4 expression. Control RAW-ASC cells primed with LPS showed increased TLR4 expression, as determined by indirect immunofluorescence using anti-TLR4 antibody and Alexa-568-labeled secondary antibody (***Fig. 2M***). In contrast, cells pretreated with Ciclopirox and Mefloquine and primed with LPS showed markedly reduced expression of TLR4 verses LPS-primed control cells (***Fig. 2M***). Interestingly, Fluoxetine did not block LPS-induced increase in TLR4 expression (***Fig. 2M***), though it did reduce basal TLR4 levels (***Fig. 2K, 2L***). These data shed some insights on how these drugs may be reducing LPS priming of NLRP3 inflammasome. To further delineate the mechanism by which these drugs may block LPS priming, we determined the status of LPS binding to the cell membrane. As shown in ***Fig. 2N***, cells treated with drugs such as Ciclopirox, Mefloquine and Fluoxetine showed a major reduction in LPS binding to the cell membrane verses control cells. However, Febuxostat-treated cells showed normal LPS binding to the cell membrane, though it blocks LPS-induced upregulation of NLRP3 and pro-IL1β. Thus, Febuxostat may be blocking LPS priming downstream of LPS-binding steps.

### Inflammasome priming blockers enhance survival in sepsis mouse model in a sex-dependent manner

To determine whether FDA-approved drugs identified as LPS-priming inhibitors confer protection *in vivo*, we evaluated their efficacy in an LPS-induced sepsis mouse model (***Fig. 3A***). We chose the drugs that showed defective LPS binding to the immune cells: Ciclopirox (30mg/kg), Mefloquine (20mg/kg), and Fluoxetine (40mg/kg) (***Fig. 3A***). Administration of a lethal dose of LPS resulted in high mortality in vehicle-treated animals (both males and females), consistent with severe systemic inflammation and septic shock (***Fig. 3B-3G***). Kaplan–Meier survival analysis demonstrated a marked delay in mortality onset and a substantial increase in overall survival rates in drug-treated cohorts (***Fig. 3B-3G***). The median survival for females in Mefloquine group (N=17) was ∼23 hours, with ****p<0.0001 by Logrank test (***Fig. 3B***), Fluoxetine group (N=9) was undefined (indicating almost complete survival after 24h) with ****p<0.0001 by Logrank test (***Fig. 3C***)., and Ciclopirox group (N=4) was∼17.7 hours vs. ∼15.7 hours for control females (N=8), *p<0.0285 by Logrank test (***Fig. 3D***). The median survival for males in Mefloquine group (N=17) was undefined with ****p<0.0001 by Logrank test (***Fig. 3E***), Fluoxetine group (N=16) showed survival rate 22.2 hours with ****p<0.0001 by Logrank test (***Fig. 3F***), and Ciclopirox (N=6) showed survival rate of undefined vs. survival rate of 14.25 hours in control males (N=11) with ****p<0.0001 by Logrank test (***Fig. 3G***). Interestingly, the effects of drugs on survival rates were sex dependent. The median survival for females in Mefloquine group was ∼23 hours (***Fig. 3B***), while males in Mefloquine group (N=17) showed higher survival, with survival rate being undefined (***Fig. 3E***). In contrast to Mefloquine, survival rate for females in Fluoxetine group was higher verses males, with females showing survival rate of undefined (***Fig. 3C***) compared to males showing survival rate of 22.2 hours (***Fig. 3F***). Females in Ciclopirox group showed survival rate of ∼17.7 hours (***Fig. 3D***), while males in Ciclopirox group showed survival rate of undefined (***Fig. 3G***).

**Fig. 3:**
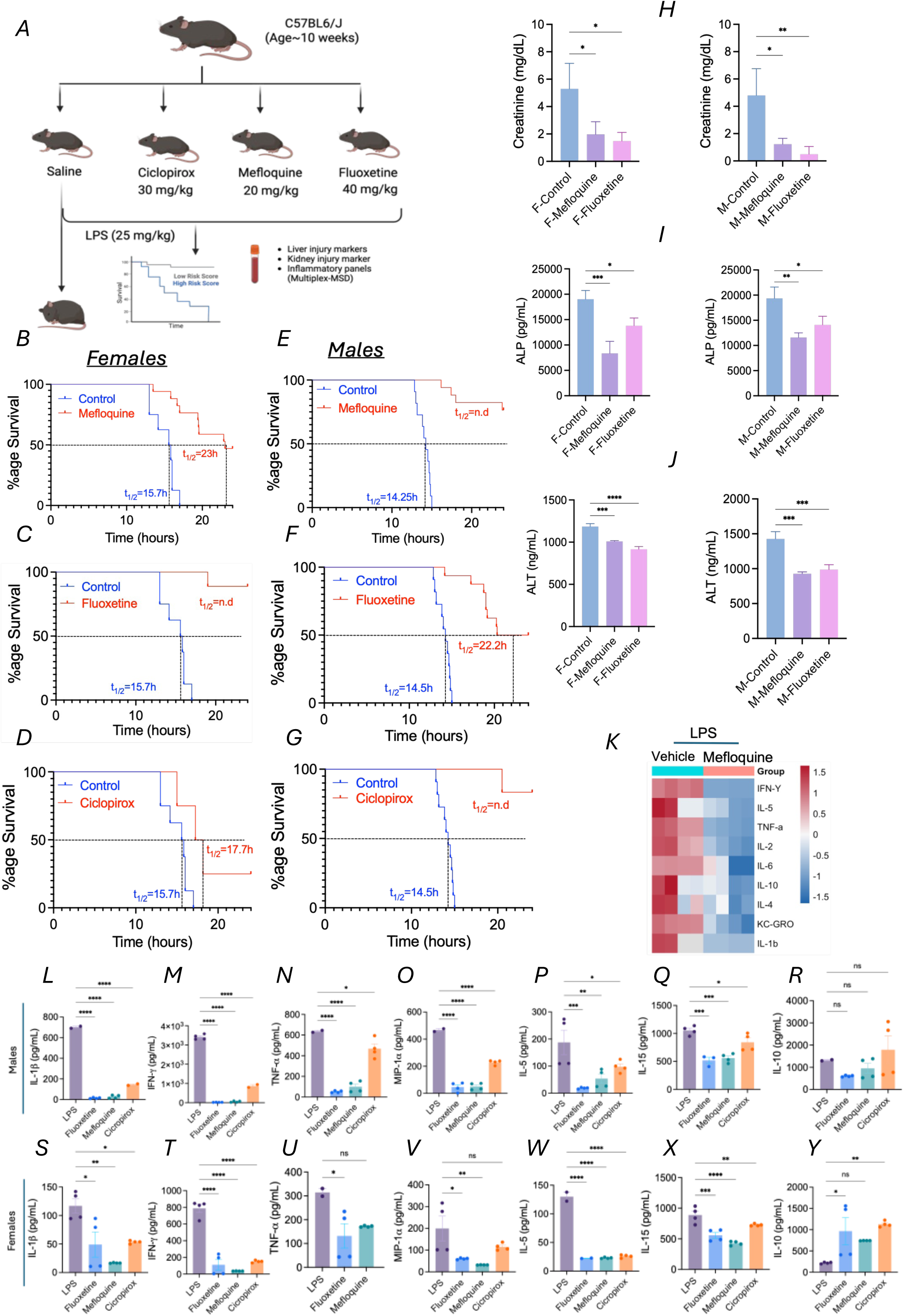
LPS priming blockers protect from sepsis-induced mortality. (***A***) Schematic diagram showing mice pretreatments with i.p injections of saline or LPS-priming blockers Ciclopirox (30mg/kg), Mefloquine (20mg/kg), and Fluoxetine (40 mg/kg). Post two hours, mice were i.p injected with a sub-lethal dose of LPS (25mg/kg). Survival curve of female mice treated with Mefloquine (***B***), Fluoxetine (***C***), or Ciclopirox (***D***). Survival curve of male mice treated with Mefloquine (***E***), Fluoxetine (***F***), or Ciclopirox (***G***). Various markers of sepsis-induced disease were determined in the serum of control or drug-treated mice and plotted for Creatinine (***H***), ALP (***I***), and ALT (***J***), with females in left row and males in right row. (***K***) Heatmap of data generated from multiplex analysis of plasma showing downregulation of various cytokines in Mefloquine-treated mice, with N=4, and scale depicts z-score with red showing increase and blue showing reduction vs. vehicle control. Multiplex analysis of plasma from male mice showing downregulation of various proinflammatory cytokines in control vs. drug-treated mice; IL-1β (***L***), IFN-γ (***M***), TNF-α (***N***), MIP-1α (***O***), IL-5 (***P***), IL-15 (***Q***), IL-10 (***R***). N=3 for each group, data plotted as mean ± SD, * indicate p <0.05 **indicate p<0.01, ***indicate p<0.001 by t test. Female mice showing downregulation of various proinflammatory cytokines in control vs. drug-treated mice; IL-1β (***S***), IFN-γ (***T***), TNF-α (***U***), MIP-1α (***V***), IL-5 (***W***), IL-15 (***X***), IL-10 (***Y***). N=3 for each group, data plotted as mean ± SD, * indicate p <0.05 **indicate p<0.01, ***indicate p<0.001 by t-test.

In addition to survival rates, we also determined if these drugs could dampen the LPS-induced damage to various organs. Compared to untreated septic mice, the mice treated with Mefloquine and Fluoxetine showed a significant reduction in kidney and liver damage, as assessed by serum levels of kidney injury marker creatinine and liver injury markers ALT and ALP. Creatinine serum levels (***Fig. 3H)*** and ALP/ALT serum levels (***Fig. 3I, 3J***) were significantly reduced in both females and males (N=6, mean±SE, ***p<0.001). To get further insights into global cytokine profile alterations, we performed unbiased multiplex analysis using the V-PLEX Mouse Cytokine 19-Plex kit and the Meso QuickPlex Q60MM instrument (Meso Scale Discovery). As shown in ***Fig. 3K***, the heatmap highlights the markedly altered cytokine profile in control verses Mefloquine male mice. Various proinflammatory cytokines, including IL-1β, TNF-1α, IFN-γ, MIP1α, IL-5, and IL-15, were significantly lower in the plasma of both male and female mice treated with Mefloquine, Fluoxetine, and Ciclopirox (***Fig. 3L-3Y***). In the same plasma sample, the levels of potent anti-inflammatory cytokine IL-10 were higher in female mice treated with Fluoxetine and Ciclopirox vs. LPS alone, while males did not show any significant effects (***Fig. 3R, Fig. 3Y***). Collectively, these findings demonstrate that pretreatment with FDA-approved drugs confer significant protection against LPS-induced lethality and organ damage, supporting their therapeutic potential for mitigating excessive inflammasome-driven inflammation in sepsis.

### Several classes of FDA-approved drugs block ASC puncta formation

To identify inhibitors of NLRP3 inflammasome assembly, THP1-ASC-GFP cells were pretreated with library drugs, primed with LPS to induce inflammasome component expression, and subsequently stimulated with the canonical NLRP3 activator Nigericin to trigger inflammasome assembly (***Fig. 4A***). Inflammasome activation was quantified using complementary readouts, including quantification of ASC speck formation, interleukin-1β (IL-1β) levels in peritoneal lavage, and multiplex cytokine array using mouse plasma. The control THP1-ASC-GFP cells showed robust ASC puncta formation, indicative of NLRP3 inflammasome assembly (***Fig. 4B***). Hits were classified based on their efficacy for blocking ASC speck formation, with reduction in % of cells showing ASC puncta ranging from 80-100%, 60-80%, 40-60% (***Fig. S2***), 20-40% (***Fig. S3***), or 10-20% (***Fig. S4***). In comparison to control (***Fig. 4D***), most potent candidate drugs included Miconazole (***Fig. 4E***), Saquinavir (***Fig. 4F***), Salmeterol (***Fig. 4G***), Rosiglitazone (***Fig. 4H***), Irbesartan (***Fig. 4I***) and Felodipine (***Fig. 4J***). To determine whether these drugs affect inflammasome assembly in murine macrophages, RAW-ASC cells were pretreated with the drugs, primed with LPS, and then activated with Nigericin to assess ASC speck formation. As shown in ***Fig. 4K-Q***, RAW-ASC cells treated with Miconazole, Saquinavir, Salmeterol, Rosiglitazone, Irbesartan, and Felodipine showed markedly reduced ASC puncta formation verses control cells treated with LPS + Nigericin. To confirm that these drugs are not affecting LPS priming or expression of NLRP3 inflammasome components, western blot analysis of inflammasome components was performed. As shown in ***Fig. 5A***, THP1-ASC-GFP cells treated with LPS or drugs + LPS showed induction in expression of NLRP3, caspase-1, and ASC-GFP (Red arrow), without affecting expression of endogenous ASC (Yellow arrow). The induced ASC-GFP expression clearly shows that Rosiglitazone, Irbesartan, and Felodipine do not alter LPS priming step (***Fig. 5A***). Cells treated with Miconazole, Saquinavir, and Salmeterol also showed LPS-induced expression of pro-IL-1β to similar levels as LPS treated THP1-ASC-GFP cells (***Fig. 5B***). RAW-ASC cells also showed that inflammasome assembly blocking drugs, such as Rosiglitazone, Irbesartan, Felodipine, Miconazole, Saquinavir, and Salmeterol do not alter the basal or LPS-induced expression of NLRP3 inflammasome components (***Fig. 5C, 5D***), indicating that inflammasome assembly is not perturbed due to lack of any inflammasome component. To evaluate whether FDA-approved drugs blocking ASC puncta formation can inhibit *in vivo* NLRP3 inflammasome assembly, we employed an LPS + ATP driven inflammasome activation model^14^. Peritoneal lavage from mice treated with LPS and ATP showed robust induction in NLRP3 inflammasome assembly, as evidenced by increased IL-1β levels in peritoneal fluid, while mice pretreated with Saquinavir markedly attenuated IL-1β levels in peritoneal lavage (***Fig. 5F)***. Similar to Saquinavir, the unbiased multiplex analysis on mouse plasma from control verses Salmeterol-treated mice showed reduced levels of proinflammatory cytokines, such as TNF-α (***Fig. 5G***), IL-β (***Fig. 5H***), IL-17A (***Fig. 5I***), IL-33 (***Fig. 5J***) and, IL-15 (***Fig. 5K***).

**Fig. 4:**
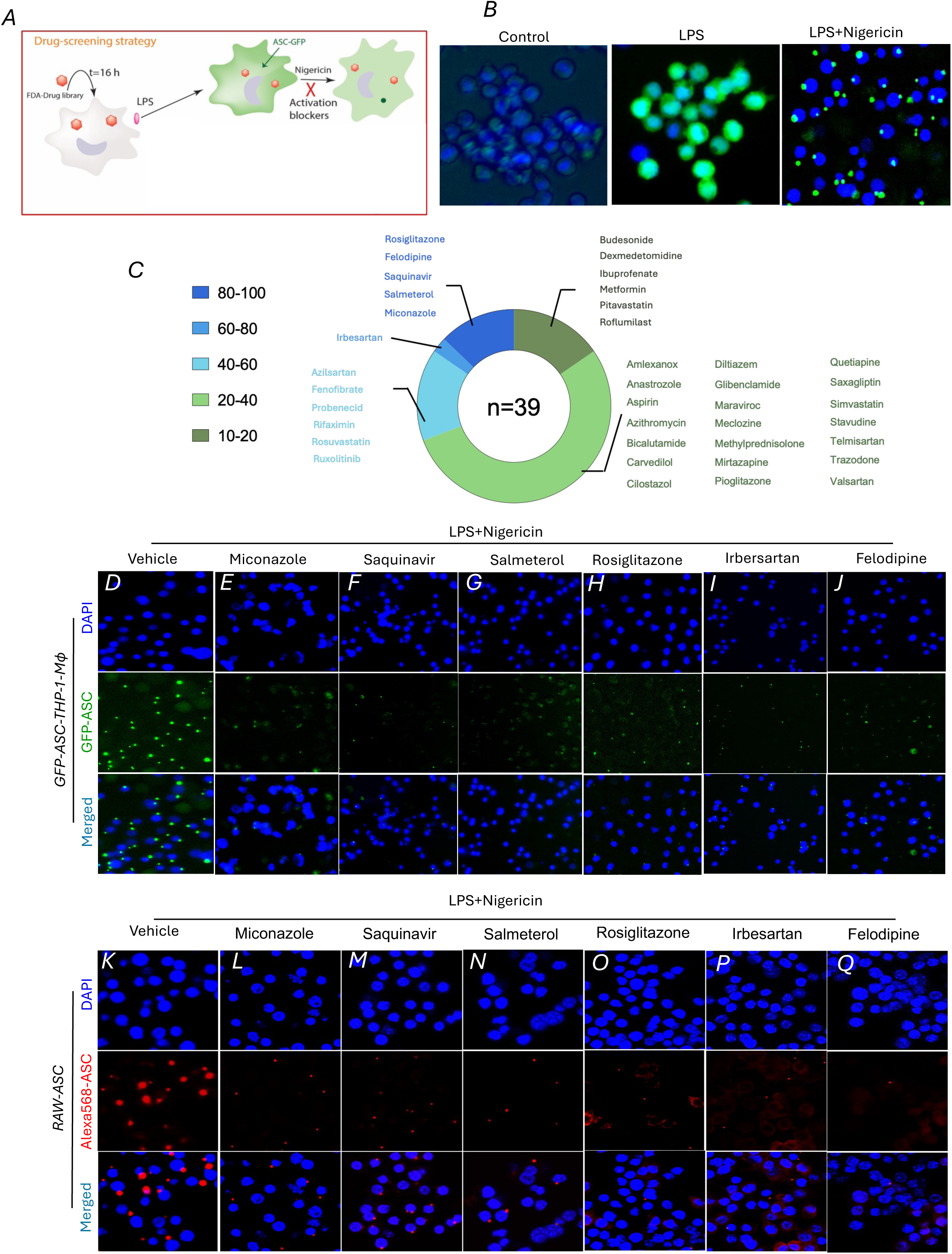
FDA-approved drugs blocking NLRP3 inflammasome assembly. (***A***) Schematic diagram showing the FDA-approved drug library screen with the human monocytic THP-1-ASC-GFP cell line. Cells were pretreated with vehicle or 20μM of the indicated drug for 16 h, followed by LPS priming and Nigericin treatment for NLRP3 inflammasome assembly (ASC puncta formation). (***B***) Control THP-1-ASC-GFP cell line stained with DAPI showing low-level basal expression of ASC-GFP. Cells treated with LPS showing induced ASC-GFP expression (central row), and cells treated with LPS+ Nigericin showing ASC-GFP puncta formation (right row). (***C***) Pie chart of 39 drugs categorized based on percentage of puncta inhibition (classified ranges include: 10-20%, 20-40%, 40-60%, 60-80%, 80-100%). THP1-ASC-GFP cells pretreated with vehicle or selected drugs for 16h, followed by washing with PBS and incubation with LPS, followed by incubation with Nigericin. ASC-puncta were visualized using fluorescent microscopy, with vehicle (***D***), Miconazole (***E***), Saquinavir (***F***), Salmeterol (***G***), Rosiglitazone (***H***), Irbesartan (***I***), and Felodipine (***J***). DAPI is used to stain nucleus. RAW-ASC cells pretreated with vehicle or selected drugs for 16h, followed by washing with PBS and incubation with LPS, followed by incubation with Nigericin. Cells were fixed and permeabilized and stained with anti-ASC antibody and Alexa-568 labeled, with images showing treatments as, vehicle (***K***), Miconazole (***L***), Saquinavir (***M***), Salmeterol (***N***), Rosiglitazone (***O***), Irbesartan (***P***), and Felodipine (***Q***). DAPI was used to stain nucleus.

**Fig. 5:**
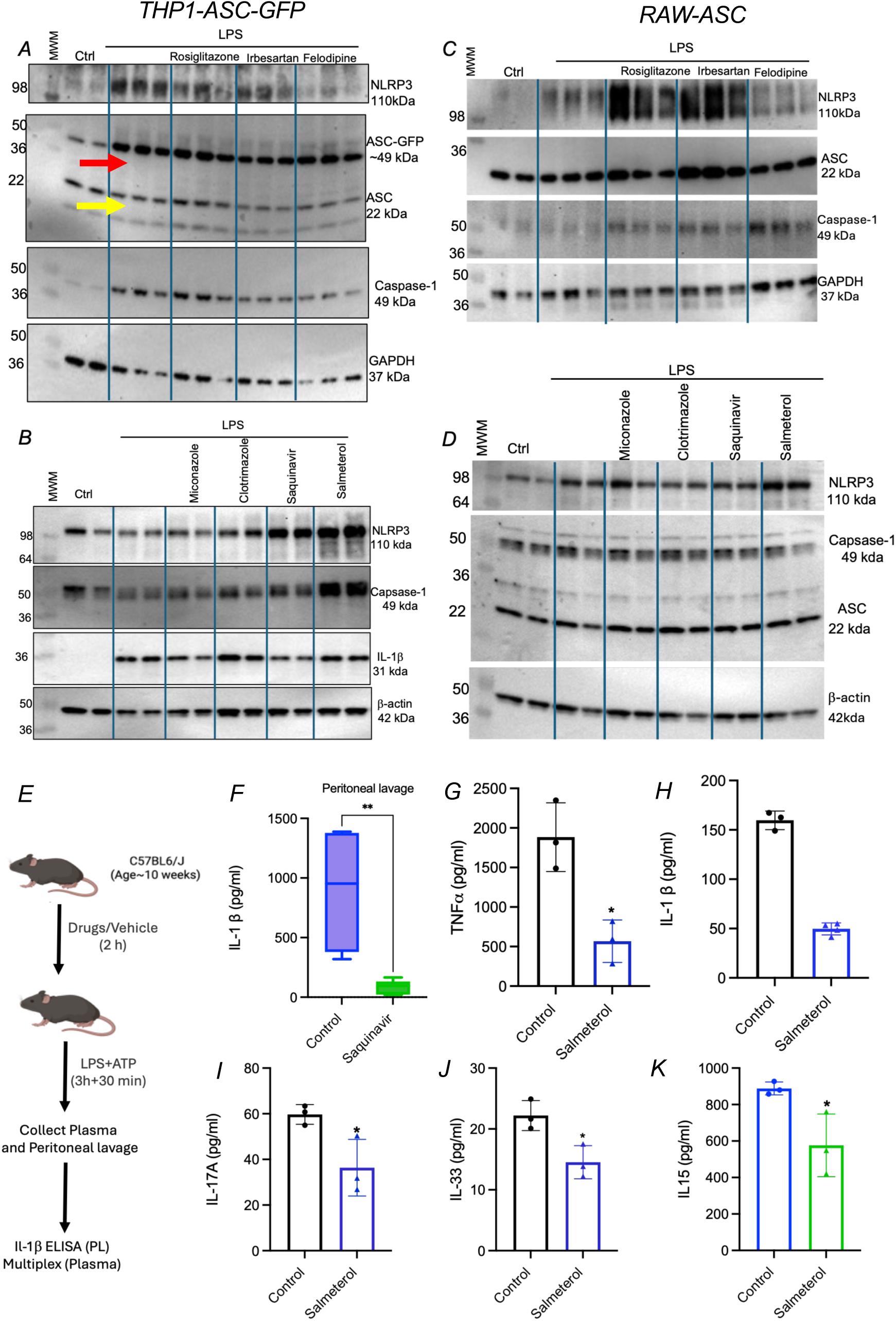
LPS priming blocking drugs reduce downstream signaling. (***A, B***) Western blot analysis of basal or LPS-induced expression of various inflammasome components in THP1-ASC-GFP cells pretreated with vehicle or selected drugs for 16h. Yellow arrow indicate endogenous ASC expression, and red arrow indicates inducible ASC-GFP expression. (***C, D***) Western blot analysis of basal or LPS-induced expression of various inflammasome components in RAW-ASC cells pretreated with vehicle or selected drugs for 16h. (***E***) Schematic of NLRP3 inflammasome assembly in vivo (***F***) Age-matched (10-week-old) male WTC57BL6J mice fed with chow diet were i.p injected with saline or saquinavir. After 2 hours, mice were primed for inflammasome assembly by an I.P. injection of LPS (5μg/mouse). After 4h of LPS injection, the NLRP3 inflammasome assembly was induced by I.P. injection of ATP (0.5 mL of 30 mM, pH 7.0). The peritoneal cavity was lavaged with 5 mL sterile PBS, and IL-1β levels in peritoneal lavage were determined by ELISA (N = 6, mean ± SD for all groups, ∗∗p < 0.01 by two-tailed t-test). Mouse plasma was used in multiplex analysis to determine levels of (***G***) TNF-α, (***H***) IL-1β, (***I***) IL-17A, (***J***) IL-33, and (***K***) IL-15 (N=3, mean ± SD for all groups, ∗p < 0.05 by two-tailed t-test).

### Effect of ASC-puncta blockers on inflammation-resolving efferocytosis and cholesterol efflux pathway

To determine whether ASC puncta blockers also enhance the inflammation-resolving functions of macrophages, we evaluated efferocytosis in macrophages. RAW-ASC cells were pretreated with Irbesartan or Rosiglitazone before incubation with fluorescently labeled apoptotic Jurkat cells. Fluorescence microscopy demonstrated that both compounds significantly increased the engulfment of apoptotic cells compared with vehicle-treated controls, with Irbesartan and Rosiglitazone showing ∼ 2-fold increase in efferocytic capacity (***Fig. 6A, 6B***). Efferocytosis is a druggable target, and efficient efferocytosis depends on the receptor tyrosine kinase MERTK, which mediates the recognition and internalization of apoptotic cells^15,16^. We therefore examined whether Irbesartan or Rosiglitazone influenced MERTK expression. Immunofluorescence analysis revealed a marked increase in total MERTK protein levels following treatment with either compound relative to vehicle controls (***Fig. 6C, 6D***), consistent with an enhanced efferocytic capacity. Because engulfment of apoptotic cells imposes a substantial cholesterol load on macrophages, successful efferocytosis requires activation of lipid-handling pathways that restore cellular homeostasis. Specifically, cholesterol derived from apoptotic cells activates the PPARγ-LXRα signaling axis, leading to induction of the cholesterol transporter ABCA1 and promoting cholesterol efflux to apolipoprotein A-1 (apoA-1). Consistent with activation of this pro-resolving pathway, Irbesartan significantly increased ABCA1 expression and enhanced cholesterol efflux to apoA-1 (***Fig. 6E, 6F***). As expected, the LXR agonist (T-compound) increased cholesterol efflux to apoA1 by ∼2-fold vs. HepG2 cells without T-compound, and Irbesartan further increased cholesterol efflux to apoA1 (***Fig. 6F***). These data indicate that Irbesartan may be modulating cholesterol efflux by enhancing ABCA1 levels, independent of LXR activation.

**Fig. 6:**
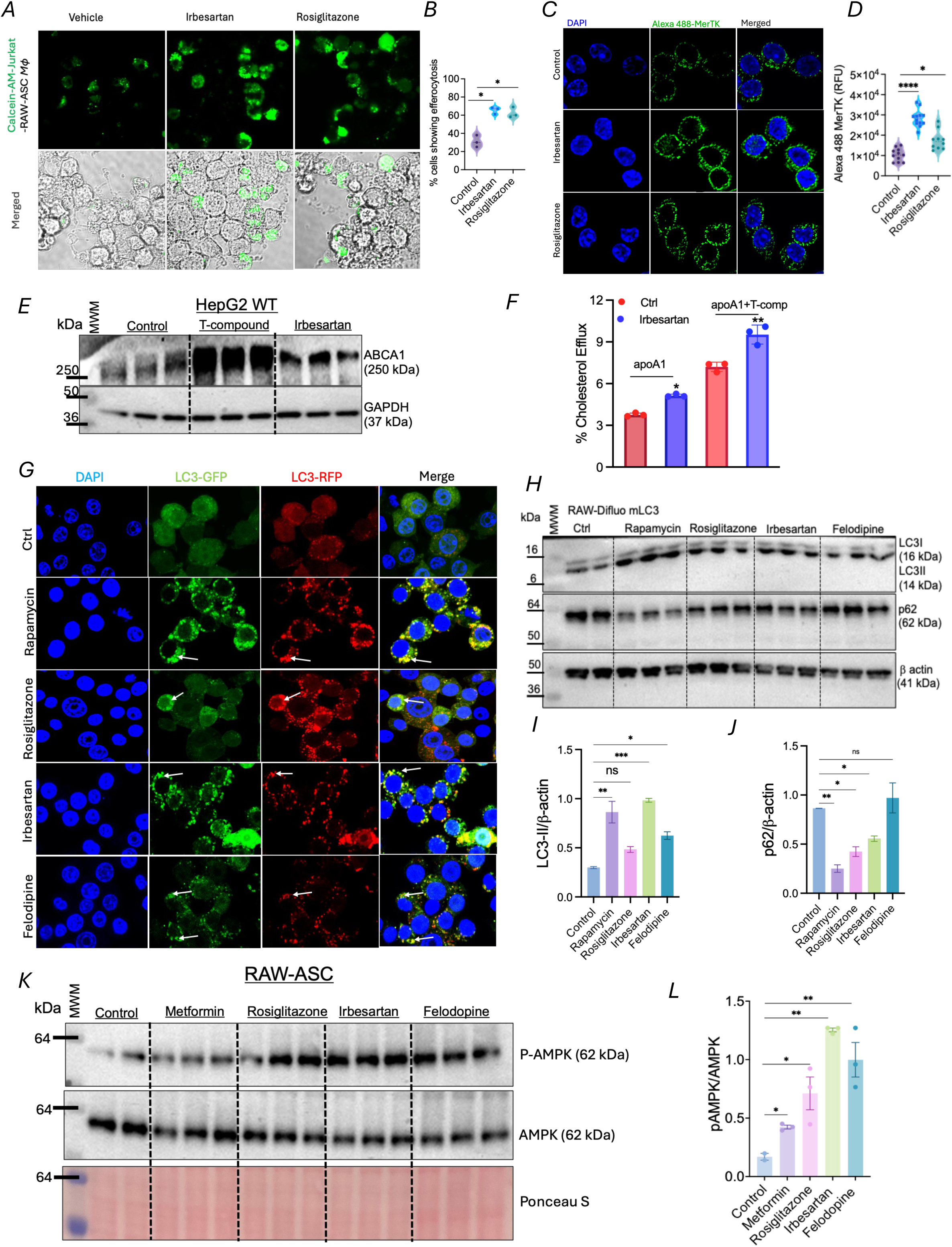
Inflammasome assembly blockers’ effect on efferocytosis, ABCA1-mediated cholesterol efflux, and autophagy. (***A***) Confocal images of RAW-ASC macrophages pretreated with vehicle, Rosiglitazone (20μM), or Irbesartan (20μM) for 16h, followed by incubation with Calcein-AM labeled apoptotic Jurkat cells for 4h. Green fluorescence represents engulfed apoptotic Jurkat cells. (***B)*** Graph showing quantification of macrophage efferocytic capacity, (N≥ 100) were analyzed. Efferocytic capacity was expressed as the percentage of macrophages containing internalized Calcein-AM positive apoptotic cells. Data are presented as mean ± SEM. Statistical significance was determined using t-test, with *p< 0.05. (***C***) Indirect immunofluorescence of MerTK receptor in non-permeabilized RAW-ASC cells treated with vehicle or Irbesartan or Rosiglitazone, using mouse anti-rabbit MerTK primary antibody and Alexa488-labeled goat anti-rabbit secondary antibody. (***D***) Graph showing quantification of MERTK levels in control vs. indicated drugs. Approximately n ≥ 10 cells were analyzed per treatment group, and relative fluorescence intensity was measured using ImageJ. Data are presented as mean ± SEM. Statistical significance was determined using One Way ANOVA (p< 0.05 versus control; ***p< 0.001 versus control). (***E***) HepG2 cells were treated with Irbesartan or T-compound (LXR agonist) for 16h, followed by western blot analysis using ABCA1 antibody. GAPDH is used as loading control. (***F***) HepG2 cells were labeled with ^3^H-cholesterol and pretreated with Irbesartan or positive control T-compound to induce ABCA1 expression. Cholesterol efflux to chase media with apoA1 ± T-compound was performed for 4 h at 37 °C in serum-free EMEM. Values are % cholesterol release, N=3, mean ± SD, ***p < 0.001 for apoA1 group, and *p < 0.05 for apoA1 + T-compound group by two-tailed t-test. (***G***) Fluorescent microscopy of RAW-Difluo mLC3 cells treated with either saline or drugs, showing LC3-GFP, LC3-RFP, and merge. (***H***) Western blot of LC3-II and p62 in RAW-Difluo mLC3 cells treated with either saline or various drugs, and (***I, J***) quantification of LC3-II and p62 vs β-actin. (***K***) Western blot analysis of total and p-AMPK in RAW-ASC cells treated with either saline or drugs. Metformin was used as positive control, and Ponceau S staining was used as loading control. (***L***) quantification of p-AMPK vs total AMPK, with N=2 for control and N=3 for all drugs, mean ± SD for all groups, * indicate p<0.05, and ** indicate p < 0.01 by two-tailed t-test.

Collectively, these findings demonstrate that ASC puncta-blocking compounds not only suppress inflammasome assembly but also enhance macrophage efferocytosis and downstream resolution-associated lipid metabolism. By increasing MERTK expression and promoting ABCA1-dependent cholesterol efflux, Irbesartan facilitates key processes required for efficient clearance of apoptotic cells and restoration of macrophage homeostasis.

### Effect of ASC-puncta blockers on inflammation-resolving autophagic pathway

To elucidate the mechanisms by which identified FDA-approved drugs suppress NLRP3 inflammasome assembly without affecting levels of inflammasome components, we examined their effects on autophagy. Autophagy is a critical regulator of inflammasome activation, particularly through the clearance of damaged mitochondria and attenuation of mitochondrial reactive oxygen species (mtROS), which serve as potent activators of NLRP3 signaling. RAW-Difluo™ mLC3 reporter cells which express tandem fluorescent LC3B to monitor autophagosome maturation were used. Treatment with Rosiglitazone, Irbesartan, or Felodipine markedly increased LC3-II RFP puncta compared with vehicle-treated controls, indicating enhanced autophagosome maturation and autolysosomal fusion (***Fig. 6G***). Consistent with these observations, immunoblot analysis demonstrated increased LC3-II accumulation accompanied by reduced p62/SQSTM1 levels, confirming enhanced autophagic flux (***Fig. 6H–6J***). Rapamycin, used as a positive control, produced comparable increases in LC3-II puncta and LC3-II protein levels together with decreased p62 expression. To further dissect, the mechanism of autophagy induction by these drugs, we determined the status of AMPK (AMP-activated protein kinase) phosphorylation. AMPK is a master metabolic sensor that promotes autophagy by directly activating the ULK1 complex and inhibiting the mTORC1 pathway. As shown in ***Fig. 6K, 6L***, Rosiglitazone, Irbesartan, and Felodipine treatment led to markedly increased phosphorylation of AMPK, with Metformin used as a positive control. The drugs that block LPS priming step, such as Fluoxetine, Mefloquine, and Ciclopirox did not show any increase in autophagic flux, rather reduced LC3-II levels and increased p62 levels, showing trend toward autophagy blockage (***Fig. S5***). Dasatinib was one of the LPS priming blocker drugs that showed robust autophagy induction (***Fig. S5***).

### Irbesartan enhances autophagy flux in mice

Given the robust induction of autophagy by Irbesartan in macrophages, we next investigated whether this effect could also be observed *in vivo*. To assess autophagic flux in live animals, we utilized the C57BL/6-(CAG-RFP/EGFP/Map1lc3b)1Hill/J transgenic mouse model, which expresses tandem fluorescent LC3 (RFP-EGFP-LC3) under the control of the CAG promoter. This reporter system enables monitoring of autophagy progression based on the differential pH sensitivity of the fluorescent proteins: both EGFP and RFP signals are detected in autophagosomes, whereas the acidic lysosomal environment quenches EGFP fluorescence while preserving RFP fluorescence following autophagosome-lysosome fusion. Consequently, a reduction in GFP signal ratio reflects increased autophagic flux. To determine the effect of Irbesartan on autophagy *in vivo*, mice were treated with Irbesartan or saline, and liver sections were analyzed for GFP-LC3 and RFP-LC3 fluorescence. Compared with controls, liver tissue from Irbesartan-treated mice exhibited a marked increase in both GFP-LC3 and RFP-LC3 puncta, indicating enhanced autophagosome formation (***Fig. 7A, 7B***). Importantly, quantitative analysis revealed a significant decrease in the GFP ratio in the Irbesartan-treated group (***Fig. 7C, D***). Because EGFP fluorescence is selectively lost following lysosomal acidification whereas RFP fluorescence remains stable, this lower GFP ratio is consistent with enhanced autophagic flux rather than an accumulation of autophagosomes due to impaired degradation. Together, these findings demonstrate that Irbesartan promotes autophagy *in vivo* and enhances autophagic flux in mouse liver.

**Fig. 7:**
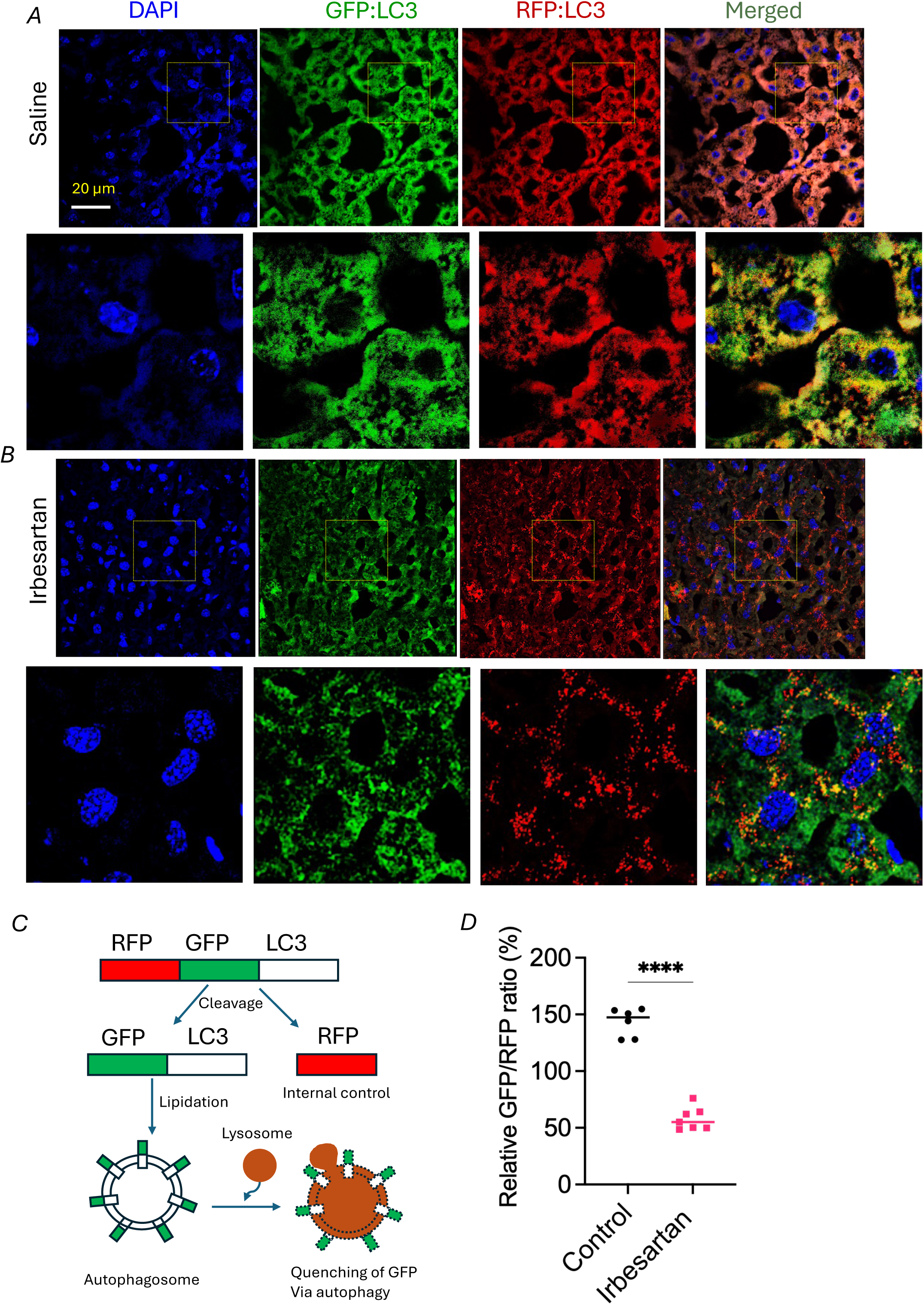
Irbesartan effect on autophagy in vivo. (***A***, ***B***) Age-matched (9-week-old) male autophagy mice fed with chow diet were i.p injected with saline or 3 mg/kg Irbesartan for 4 days. After 4 days, the liver was harvested and OCT embedded. Liver tissue was sectioned and confocal microscopy was performed. DAPI staining is used to visualize nuclei. (***C***) Schematic showing autophagy induction and flux in autophagy reporter mouse, and (***D***) Quantification of relative GFP/RFP percentage ratio. N=3 mice, 6 sections were analyzed, mean ± SEM for all groups, **** indicate p<0.0001 by t-test.

### Effect of ASC-puncta blockers on mitochondrial dynamics, mitochondrial ROS, and mitochondrial membrane potential

LPS promotes Drp1-dependent mitochondrial fission and associated inflammatory responses in macrophages^17,18^. To determine if Irbesartan preserves mitochondrial integrity, we examined mitochondrial morphology in differentiated THP-1 macrophages following LPS stimulation using the mitochondrial outer membrane marker TOM20. In vehicle-treated macrophages, LPS priming induced profound mitochondrial remodeling characterized by fragmentation of the normally interconnected mitochondrial network, showing a marked increase in punctiform mitochondria accompanied by a loss of elongated tubular structures (***Fig. 8A***), indicating mitochondrial hyper-fragmentation. In contrast, macrophages pretreated with Irbesartan retained an extensive mitochondrial network despite subsequent LPS exposure. Enlarged regions (orange boxes) demonstrated preservation of elongated mitochondria and a substantial reduction in fragmented punctiform structures. Representative examples and quantifications of elongated (E), intermediate (I), and punctiform (P) mitochondria are indicated in the insets (***Fig. 8A, 8B***).

**Fig. 8:**
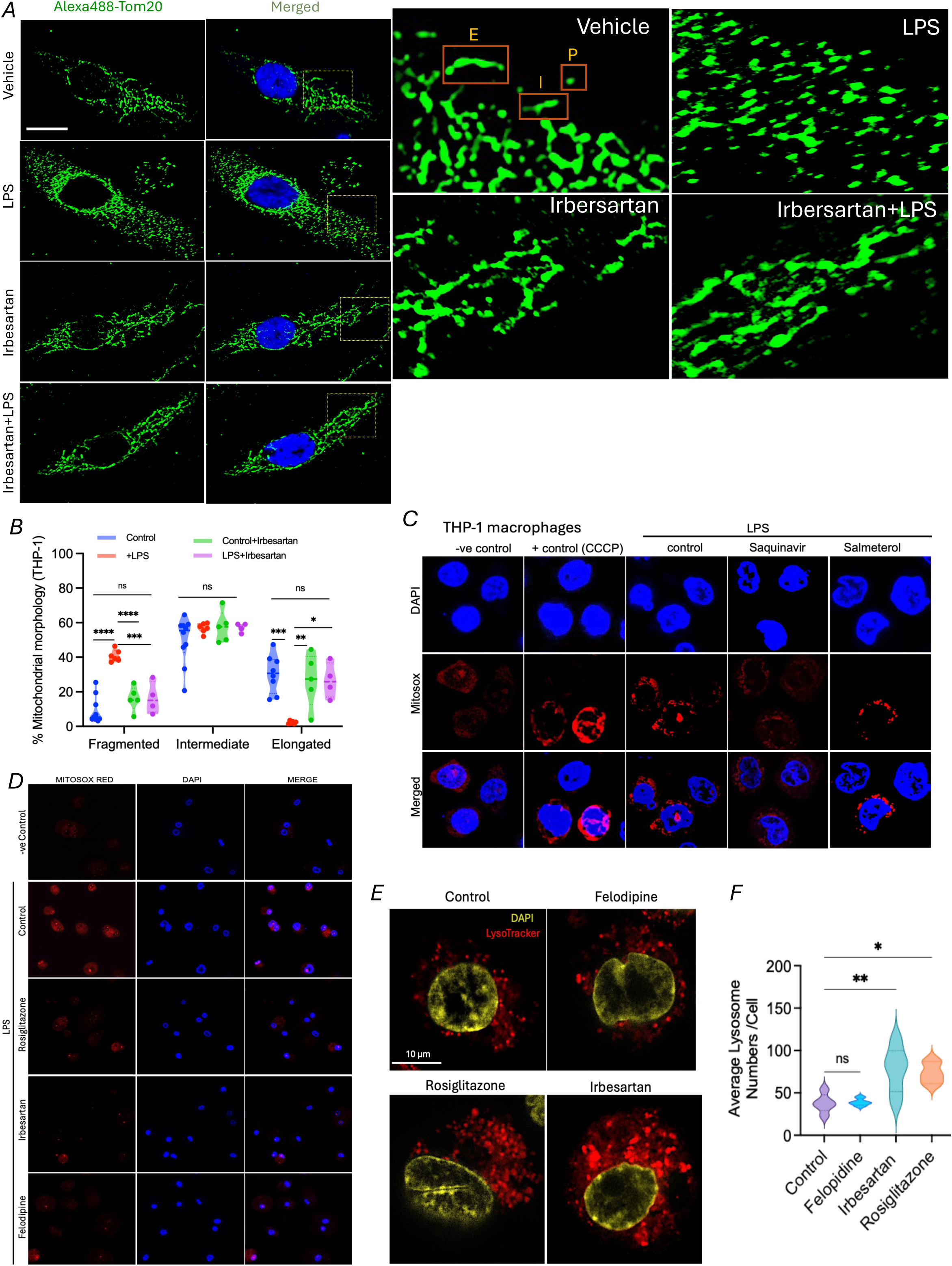
ASC-puncta blockers attenuate LPS-induced mitochondrial fragmentation and mROS and increase lysosomal biogenesis. (***A***) Confocal immunofluorescence showing Tom20-labeled mitochondria in differentiated THP-1 macrophages treated vehicle, LPS, Irbesartan, or LPS + Irbesartan, with right panel showing magnified image from yellow boxed region. Regions demarcated by orange boxes are examples of elongated “E,” intermediate “I,” and punctiform “P” mitochondria, as marked by letters. (***B***) Classification of mitochondrial morphology based on Mitoprofiler (ImageJ), with N≥ 10 and plotted values are mean ± SEM for each type of mitochondrial, ns=non-significant, * indicate p <0.05 **indicate p<0.01, ***indicate p<0.001 and ****indicate p<0.0001 by t-test. (***C, D***) THP-1 macrophages pretreated with vehicle or selected drugs for 16h, followed by washing with PBS and incubation with LPS (1µg/ml) for 3 hours, followed by staining with Mitosox Red dye and fluorescent microscopy. DAPI is used to visualize nuclei. (***E***) Stimulated Emission Depletion Super-Resolution Imaging microscopy (STEDYCON) of THP-1 macrophages treated with vehicle, Felodipine, Rosiglitazone, and Irbesartan showing lysosomes labeled with lysotracker. (***F***) Quantification of lysosome number per cell and the values are plotted as mean ± SEM, where * indicate p <0.05, **indicate p<0.01, and ns is non-significant by t-test.

Given the established role of mitochondrial dysfunction in NLRP3 activation, we next assessed the impact of ASC-puncta blockers on LPS-induced mtROS production. Carbonyl cyanide m-chlorophenyl hydrazone (CCCP) was used a positive control, as it induces mitochondrial depolarization and stimulate mitoROS production by acting as a protonophore uncoupler of oxidative phosphorylation (***Fig. 8C***). LPS priming resulted in a significant increase in mitochondrial reactive oxygen species (mtROS) levels while treatment with Saquinavir and Salmeterol markedly reduced mtROS accumulation compared with LPS alone (***Fig. 8C***), indicating improved mitochondrial redox homeostasis. Similar results were found with Irbesartan, Felodipine, and Rosiglitazone, with all showing clear reduction in mtROS levels (Fig***. 8D***). This reduction in mtROS supports a functional link between mitochondrial ROS suppression and impaired inflammasome priming and activation.

As lysosomal leakage, either induced by cholesterol crystal or other particulate matter, plays a major role in inflammasome activation^19,20^, we tested if Felodipine, Rosiglitazone, or Irbesartan modulated the lysosomal number. THP-1 macrophages were treated with and analyzed by LysoTracker staining using high-resolution STEDYCON super-resolution microscopy (***Fig. 8E, 8F***). LysoTracker fluorescence demonstrated a marked increase in the abundance of acidic lysosomal vesicles in cells treated with Irbesartan and Rosiglitazone compared with vehicle-treated controls, while Felodipine showed no effect. Quantitative image analysis confirmed a significant increase in lysosomal number per cell following treatment with Rosiglitazone and Irbesartan, indicating that this compound enhances lysosomal biogenesis and/or lysosomal accumulation (***Fig. 8E, 8F***). The increase in lysosomal content is consistent with activation of cellular degradative pathways and suggests that these drugs promote clearance of inflammatory substrates and damaged organelles.

## Discussion

Dysregulated NLRP3 activity is involved in numerous chronic inflammatory and metabolic disorders. The final executioner of inflammasome activity, “GSDMD” promotes LPS-induced sepsis and mortality^6,8^. Several studies, including those from our lab, have shown that GSDMD can be potently blocked by an FDA-approved drug, Disulfiram (DSF), and that DSF can serve as a therapeutic option for a variety of diseases, such as obesity and atherosclerosis^21–25^. Our findings reveal that multiple agents traditionally used for unrelated indications possess significant inhibitory activity against NLRP3 inflammasome priming as well as activation. The identified drugs belong to various classes of drugs based on their actions, such as antidepressants, antimalarials, antifungals, antivirals, anti-COPD, and anti-CVD agents.

The priming blockers include drugs from various categories. For example, Febuxostat is an FDA-approved xanthine oxidase inhibitor used clinically for treating gout. By inhibiting uric acid production, Febuxostat reduces monosodium urate crystal formation, a key trigger of NLRP3 inflammasome activation in gout^26^. Febuxostat also exert anti-inflammatory effects by attenuating release of pro-inflammatory cytokines such as IL-1β^27^. Our data show that Febuxostat blocks LPS priming, providing new mechanistic insights into the anti-inflammasome activity of this drug. Dasatinib was shown to suppress particulate-induced pyroptosis and acute lung inflammation^28^, while Ciclopirox was shown to block inflammasome activation^29^. Dasatinib is a broad-spectrum tyrosine kinase inhibitor approved for the treatment of chronic myeloid leukemia. Similarly, Tamoxifen is a hormone therapy medication used to treat and prevent hormone receptor-positive breast cancer. Our data show that Dasatinib and Tamoxifen block LPS priming, which may limit the activity of tumor-promoting immune cells such as tumor-associated macrophages (TAMs) and myeloid-derived suppressor cells (MDSCs). Mefloquine is an antimalarial drug approved for Plasmodium infections. Beyond its antiparasitic activity, Mefloquine has been shown to modulate innate immune responses by altering lysosomal function and disrupting endolysosomal acidification. These processes are increasingly recognized as regulators of inflammasome signaling. Mefloquine has been reported to suppress pro-inflammatory cytokine production and inhibit inflammasome activation^30^. Our data showing that Mefloquine reduces LPS priming via blocking LPS binding to the cell membrane provide new mechanistic insights into the anti-inflammasome activity of this drug. Further, Mefloquine reduced plasma levels of pro-inflammatory cytokines in both sexes but provided more robust protection against sepsis in males vs. females, indicating sex-specific effects (***Fig. 3***). Interestingly, a previous study showed that plasma Mefloquine concentrations after 14 hours were considerably higher in human female subjects than in males^31^. Fluoxetine (Prozac) is a selective serotonin reuptake inhibitor widely prescribed for the treatment of major depressive disorder and anxiety-related conditions. An elegant recent study showed that Fluoxetine protects from sepsis via promoting IL-10-dependent metabolic defenses^32^. Fluoxetine has also been shown to inhibit NF-κB activation, reduce expression of pro-inflammatory cytokines, and suppress microglial and macrophage activation^33^. Our data show that Fluoxetine reduced LPS binding to the cell surface of immune cells, providing additional mechanistic insights into the mechanisms regulating anti-inflammasome activity of Fluoxetine. Fluoxetine effects on LPS-induced septic mortality seem sex-dependent, as females showed much higher survival rates than males. Our multiplex analysis showed that most proinflammatory cytokines were reduced in both males and females treated with Fluoxetine, while levels of the potent anti-inflammatory cytokine IL-10 were higher only in females. Ciclopirox is an FDA-approved antifungal agent and its ability to modulate cellular redox balance and inflammatory transcriptional programs suggests a potential role in suppressing priming in innate immune cells, and our data showing the direct effect of Ciclopirox on LPS binding to the cell membrane corroborate this notion. Males treated with Ciclopirox showed much higher survival rates vs. females. Interestingly, a recent study using male mice showed that Ciclopirox protects from sepsis via inactivation of SORT1-mediated wnt/β-Catenin signaling pathway^34^. FTY720 (fingolimod) is an FDA-approved sphingosine-1-phosphate (S1P) receptor modulator used for the treatment of relapsing forms of multiple sclerosis^35^. Fingolimod exerts potent immunosuppressive effects by sequestering lymphocytes in lymphoid organs. Our data corroborate these findings by showing FTY720 can also block priming of immune cells. Amlodipine, a functional inhibitor of acid sphingomyelinase, is a calcium channel blocker used to treat high blood pressure, heart disease, and angina. Amlodipine was recently shown to be associated with a lower risk of acute outpatient infections, including sepsis^36^. Our data showing that Amlodipine can block LPS priming in immune cells provided additional mechanistic insights on Amlodipine’s mechanism of action. Several antiviral and antibacterial drugs were also identified as inflammasome blockers, corroborating earlier studies^37^.

Next, we focused on drugs that can block NLRP3 inflammasome assembly, without altering basal or LPS-induced expression of inflammasome components. We categorized the selected drugs based on their efficacy in blocking NLRP3 inflammasome assembly. Several potent candidates (showing 80-100% reduction in ASC puncta) were identified, such as, Rosiglitazone, Irbesartan, Saquinavir, Salmeterol, Miconazole, and Felodipine. Recent studies showed that Miconazole attenuates LPS-induced lung inflammation by modulating alveolar macrophage polarization^38^ and attenuates neuroinflammation through the suppression of NF-kB activation and iNOS production^39^. Previous investigations demonstrated that Saquinavir, a first-generation HIV protease inhibitor, inhibits TLR4-mediated signaling^40^. The long-acting β2-adrenergic receptor agonist Salmeterol exerted anti-inflammatory effects in the pulmonary compartment of humans exposed to LPS^41^. Rosiglitazone is used to treat type 2 diabetes, while Irbesartan reduces blood pressure, protects kidney function in type 2 diabetes patients, and is also used to manage heart failure. Felodipine, a calcium channel blocker, has been shown to decrease production of proinflammatory cytokines while reducing oxidative stress and tissue injury in neurodegenerative disease^42^. Our data showing that these drugs block assembly of NLRP3 inflammasome provide new mechanistic insights into the mechanism of the action of these drugs on dampening inflammation. Furthermore, our data showing that Rosiglitazone, Irbesartan, or Felodipine can robustly induce efferocytosis and autophagy in macrophages via the AMPK pathway, provides new mechanisms for anti-inflammatory activity of these drugs. Efferocytosis plays a major role in human health and disease^15^. In addition to removal of dead cells, macrophages need to efflux the engulfed cholesterol to liver, as cholesterol crystals can activate NLRP3 inflammasome^43,44^. Irbesartan induces ABCA1 expression and cholesterol efflux, providing additional mechanism for blocking inflammasome activation. Autophagy is a critical regulatory mechanism governing innate immune signaling, particularly in the control of NLRP3 inflammasome assembly and activation in heart disease^45,46^. One of the most well-established mechanisms by which autophagy restrains NLRP3 inflammasome assembly is through the removal of damaged mitochondria via mitophagy^47^. Mitochondrial dysfunction is a potent trigger of NLRP3 activation, primarily through the generation of mitochondrial reactive oxygen species (mtROS), release of oxidized mitochondrial DNA (mtDNA), and perturbations in mitochondrial membrane potential. Thus, our data showing that Rosiglitazone, Felodipine, and Irbesartan induce autophagy via phosphorylation of AMPK adds new mechanistic details about anti-inflammasome activity of these drugs. Disease-associated defects in autophagy, such as those observed in aging, metabolic syndrome, and neurodegeneration, may predispose tissues to heightened NLRP3 inflammasome activation, thus the drugs identified in our screen can be repurposed for broad spectrum of diseases. Nevertheless, there are some inherent limitations of *in vitro* screening. The screen was conducted under defined inflammatory stimuli that may not capture the full spectrum of physiologic NLRP3 activation.

In conclusion, leveraging the Tocriscreen FDA-approved library has enabled the rapid identification of clinically relevant drugs with anti-inflammasome activity. These findings provide a foundation for both mechanistic exploration and translational development, with the aim of expanding therapeutic options for diseases promoted by dysregulated inflammasome activity.

## Materials and Methods

### Cell Culture and Reagents

All mammalian cells were maintained at 37°C with 5% CO2. The RAW264.7 cells (ATCC), RAW264.7-ASC (InvivoGen), RAW-Difluo™ mLC3 (InvivoGen), THP-1 (ATCC), THP1-ASC-GFP (InvivoGen), HepG2 WT (ATCC) were cultured in appropriate media containing required growth factors and antibiotics. RAW-ASC (Invivogen #raw-asc) were maintained in DMEM (Cleveland Clinic Media Core #11-500p) supplemented with 10% FBS, 1% Pen/Strep 5000u/mL, 100 µg/ml Blasticidin (Invivogen #ant-bl) and 100 µg/ml Normocin (Invivogen #ant-nr) RAW-Difluo™ mLC3 (Invivogen # awdf-mlc3) were maintained in DMEM supplemented with 10% FBS (Gibco), 1% pen-strep, 100 µg/ml Zeocin (Invivogen #ant-zn) and 100 µg/ml Normocin (Invivogen). THP1-ASC-GFP cells (Invivogen #thp-ascgfp) were maintained in RPMI 1640, 2 mM L-glutamine, 25 mM HEPES, 10% heat-inactivated fetal bovine serum, 100 μg/ml Normocin™,1% Pen/Strep. THP-1 cells (ATCC # TIB 202) were maintained in RPMI 1640 (Cleveland Clinic Media Core #10-500p) containing 10% FBS, 1% Pen/Strep, 0.05 mM 2-mercaptoethanol (Sigma #M3148). THP1 cells were differentiated into macrophages using 100ng/mL phorbol 12-myristate 13-acetate (Sigma P8139) for 3 days.

### Mouse models for NLRP3 inflammasome assembly

The C57BL6J-WT mice were purchased from Jackson Laboratories. All animal experiments were performed in both sexes and in accordance with approved protocol from the Cleveland State University Institutional Animal Care and Use Committee (IACUC). Mice were maintained in a temperature-controlled facility with a standard 12-hour light/dark cycle, with ad libitum food/water. For in vivo NLRP3 inflammasome study, the chow-fed WT mice were i.p injected with either vehicle, saquinavir(10mg/kg), or salmeterol (0.16 mg/kg). After 2 hours, mice were i.p injected with either saline or 5μg LPS (Escherichia coli 055:B5, Sigma) for 4h. The Nlrp3 inflammasome assembly in mice was induced by i.p injection of ATP (0.5 ml of 30 mM, pH 7.0), as described in our earlier study^14^. The mice were euthanized after 30 min, and the peritoneal cavity was lavaged with 5 ml PBS. Approximately 3.5 ml peritoneal lavage fluid was recovered from each mouse and centrifuged at 15K rpm for 10 min at room temperature. The supernatant was subjected to IL-1β ELISA, using mouse IL-1β Quantikine ELISA kit (R&D Systems # MLB00C) following the manufacturer’s instructions.

### Mouse model for LPS-induced sepsis

For LPS-induced sepsis mouse model, mice were randomly divided into saline, Ciclopirox (30 mg/kg), Mefloquine (20 mg/kg), or Fluoxetine (40 mg/kg). After 2 hours, a lethal dose of LPS (25 mg/kg) was injected i.p to induce the endotoxemia. The survival rate was determined using a log-rank statistical approach. The plasma from these mice was used for multiplex cytokine array assay using V-PLEX Mouse Cytokine 19-Plex kit and Meso QuickPlex SQ120 (Meso Scale Discovery). Serum ALT (Abcam), ALP (Abcam), and creatinine (Crystal Chem) were measured by assay kits following manufacturer’s instructions.

### LPS priming and ASC speck formation assay in THP-1-ASC-GFP cells

THP1-ASC-GFP cells were resuspended at 2 x 10^6^ cells/ml and were plated in 96-well plates with dilution of ∼ 3 x 10^5^ cells/per well. Cells were primed by treatment with LPS (1 ug/ml) for 4h at 37 °C in 5% CO2. The priming was assessed microscopically by visualization of the intracellular GFP signal. After confirming priming, the priming medium was removed and fresh media containing inflammasome inducer Nigericin (5-10 μM) was added to primed cells. Cells were incubated for 1 hour at 37 °C in 5% CO2, and fluorescent ASC specks were monitored in real-time using confocal fluorescence microscopy using normal FITC filter sets and imaged using the Nikon Eclipse Ti confocal and Nikon NIS Elements Imaging Software version 4.1.

### Autophagy Reporter Assay

RAW-Difluo mLC3 were plated in chamber slides (Ibidi; #80427). Cells were treated with ± 20 µM selected drugs for 16h, then washed with 3x with PBS, fixed in 3.7% paraformaldehyde for 3m, washed with PBS, and then counterstained for DAPI (Invitrogen) and mounted. Microscopy was performed using the Nikon Eclipse Ti confocal and Nikon NIS Elements Imaging Software version 4.1.

### Autophagy Mouse Model

The mouse model C57BL/6-Tg-(CAG-RFP/EGFP/Map1lc3b)1Hill/J was used for studying the effect of drugs on in vivo autophagic flux. The CAG-RFP-EGFP-LC3 transgenic mice have CAG promoter/enhancer sequences driving expression of a red fluorescent protein (RFP), an enhanced green fluorescent protein (EGFP), and a microtubule-associated protein 1 light chain 3 alpha (Map1lc3a or LC3) gene. The transgenic mice express RFP and EGFP in a pH-dependent manner in phagocytic cells upon autophagy induction and progression. Male mice (9 weeks old) were injected i.p with 3mg/kg Irbesartan (in 0.9% normal saline) for 4 consecutive days. Combined GFP and RFP fluorescence yields a yellow signal within high-pH phagophores and autophagosomes, while EGFP is quenched in autolysosomes, and they emit only an RFP signal.

### Efferocytosis

RAW-ASC macrophages were seeded onto 8-well ibidi chamber slides and allowed to adhere overnight. Cells were treated with the indicated drugs (20 μM) for 16 h. Jurkat T cells (ATCC, catalog number) were labeled with 1 μM Calcein-AM for 1 h at 37°C, followed by three washes with phosphate-buffered saline (PBS) to remove unbound dye. Subsequently, labelled Jurkat cells were treated with 1 μM Staurosporine for 16 h to induce apoptosis. Apoptotic Jurkat cells were co-incubated with RAW-ASC macrophages at a 1:1 ratio for 4 h at 37°C. Following incubation, non-engulfed apoptotic cells were removed by washing three times with PBS (5 min each). Cells were fixed with 3.7% paraformaldehyde for 30 min at room temperature, washed three times with PBS. Cells were imaged using confocal laser scanning microscopy using FITC-488 laser and DIC. Efferocytosis was quantified as the percentage of macrophages containing internalized Calcein-AM positive apoptotic Jurkat cells.

### MerTK expression

RAW-ASC macrophages were treated with the indicated compounds for 16 h and fixed with 3.7% methanol-free paraformaldehyde for 30 min at room temperature. Cells were washed three times with ice-cold PBS and blocked with 5% normal goat serum prepared in 1% bovine serum albumin (BSA) in PBS. Cells were incubated with anti-MerTK primary antibody (Thermo Fisher Scientific, PA5-15028) for 2 h at room temperature or overnight at 4°C. Following primary antibody incubation, cells were washed three times with PBST (0.05% Tween-20 in PBS) for 5 min each and incubated with Alexa Fluor-conjugated secondary antibodies (Invitrogen; dilution 1:1000) for 1 h at room temperature in the dark. Cells were washed three additional times with PBST, counterstained with DAPI, and mounted for confocal microscopy. MerTK fluorescence intensity was quantified using ImageJ software and expressed as relative fluorescence intensity per cell.

### Cholesterol efflux assay

The cholesterol efflux assays were performed in HepG2 cells. The cells were plated in 24-well plates at a density of 300-400,000 cells per well. The cells were labeled with ^3^H cholesterol in 1% FBS in EMEM for 24h. The labeled cells were induced for ABCA1 expression for 16h with LXR agonist T-compound (0.5 μM). Cells were washed with serum free media and chased for 4h in serum-free EMEM in the presence or absence of apoA1 (5 μg/ml). The radioactivity in the chase media was determined after brief centrifugation to pellet any residual debris. Radioactivity in the cells was determined by extraction in hexane: isopropanol (3:2) with the solvent evaporated in a scintillation vial prior to counting. The percent cholesterol efflux was calculated as 100 × (medium dpm) / (medium dpm + cell dpm).

### Western Blotting

Western blot analysis was performed on protein extracts prepared by using MPER lysis buffer + protease inhibitors (Sigma P8340) + PMSF (Sigma P7626) ± various treatment conditions. Equal amounts of proteins (∼40-50 μg/ml) from cell extracts, determined by using BCA assay (Pierce) or Nanodrop 2000 (Thermo Fisher Scientific) were resolved on a Novex 4-20% Tris-Glycine Gel (Invitrogen) then transferred to a PVDF membrane (Invitrogen). After blocking in Casein (Thermo Fisher Scientific#37525), the blots were probed with respective primary antibodies 1:1000 for overnight, washed with PBS + Tween 20 (Fisher Chemical; #BP337-500) then probed with a 1:10,000 dilution of respective secondary antibody for 1 hour, then washed with PBST. Blots were developed using the Immobilon Western Chemiluminescent HRP Substrate (Millipore; #WBKLS05000 and captured using the iBright™ CL750 Imaging System (Invitrogen; #A44116). Blots were probed with antibodies from Cell Signaling (NLRP3, β-actin, GAPDH, LC3-II, p62, ASC, IL-1β, AMPK, P-AMPK), Caspase-1 (Thermo Fisher Scientific), or ABCA1 (Novus Biologicals).

### Indirect Immunofluorescence

RAW-ASC cells were grown in regular growth media. THP-1 cells were grown in chamber slides and differentiated for 72 hours using 100ng/ml of PMA (Sigma P8139). On the 3rd day, treatment is done as indicated and cells were fixed using 3.7% methanol-free paraformaldehyde for 30 minutes. After 30 minutes, cells were washed with ice-cold PBS 3 times and permeabilized using 0.2% triton-X for 10 minutes at RT. After permeabilization, cells were washed with PBS and then subjected to blocking using 5% normal Goat serum in 1% BSA in PBS, followed by incubation with primary antibodies against NF-kB or ASC antibodies for 2 hours at RT or overnight at 4°C.

### IL-1β ELISA

THP-1 cells were treated with LPS (1μg/ml) for 3 hours and with Nigericin (5μM) for 1 hour. After treatment, media was collected and spun at 12000 RPM for 5 minutes to collect supernatant. Plasma samples from mice were used directly. IL-1β was measured using an IL-1β ELISA kit specific for humans (R&D system # DLB50) or mice (R&D system # MLB00C).

### Mitochondrial morphology

Confocal immunofluorescence imaging of the mitochondrial marker Tom20 in THP-1 macrophage were performed to analyze mitochondrial morphology. Briefly, cells were grown in 4-chambered slide and treated with 20 µM of Irbesartan and DMSO as control for 16 hours. Subsequently, cells were washed with PBS three times and 1 µg/mL of LPS was added to the cells for 3 hours for priming signal of inflammasome assembly. Next, we fixed the cells with 3.7% paraformaldehyde for 15 minutes and washed with cold PBS three times in 5-minute interval. Cells were permeabilized with 0.1% Triton-X for 10 minutes and washed and blocked with 5% goat serum at room temperature. Cells were incubated overnight at 4°C against Tom20 (1:2000; Cell Signaling: D8T4N# 42406). Followed by Alexa Flour 488-labeled secondary antibodies (2 h at room temperature). After staining of nuclei with Hoechst dye (1:10,000 dilution), cells were mounted onto slides and imaged using an Olympus confocal microscope. The predominant mitochondrial morphology in expressing cells was used for classification for elongated, intermediate and fragmented mitochondria. The classification was done by MitoProfiler plugin in ImageJ.

### Mitochondrial ROS

Mitochondrial ROS levels were determined in control or drug-treated THP-1 macrophages (PMA-treated for 72h). Cells were primed with 1ug/ml LPS for 4h, followed by staining with MitoSOX™ Red Mitochondrial Superoxide Indicator (Thermo Fisher Scientific) using manufacturer’s protocol.

## Supplementary Information

**Fig. S1:**
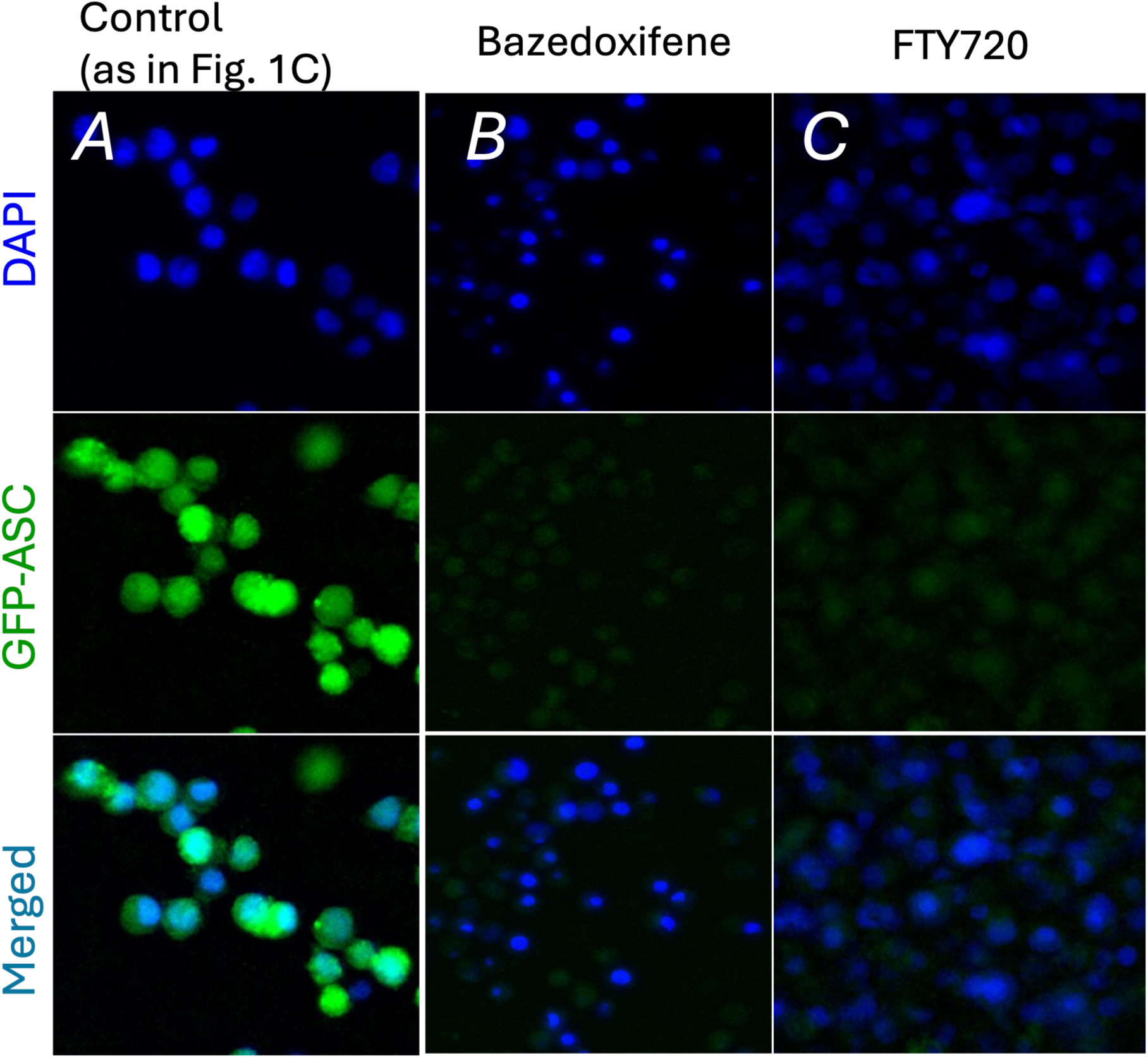
Screening of FDA-approved drugs identifies drugs blocking LPS priming. (*A)* Bazedoxifene (*B)* Bazedoxifene (*C)* FTY720

**Fig. S2:**
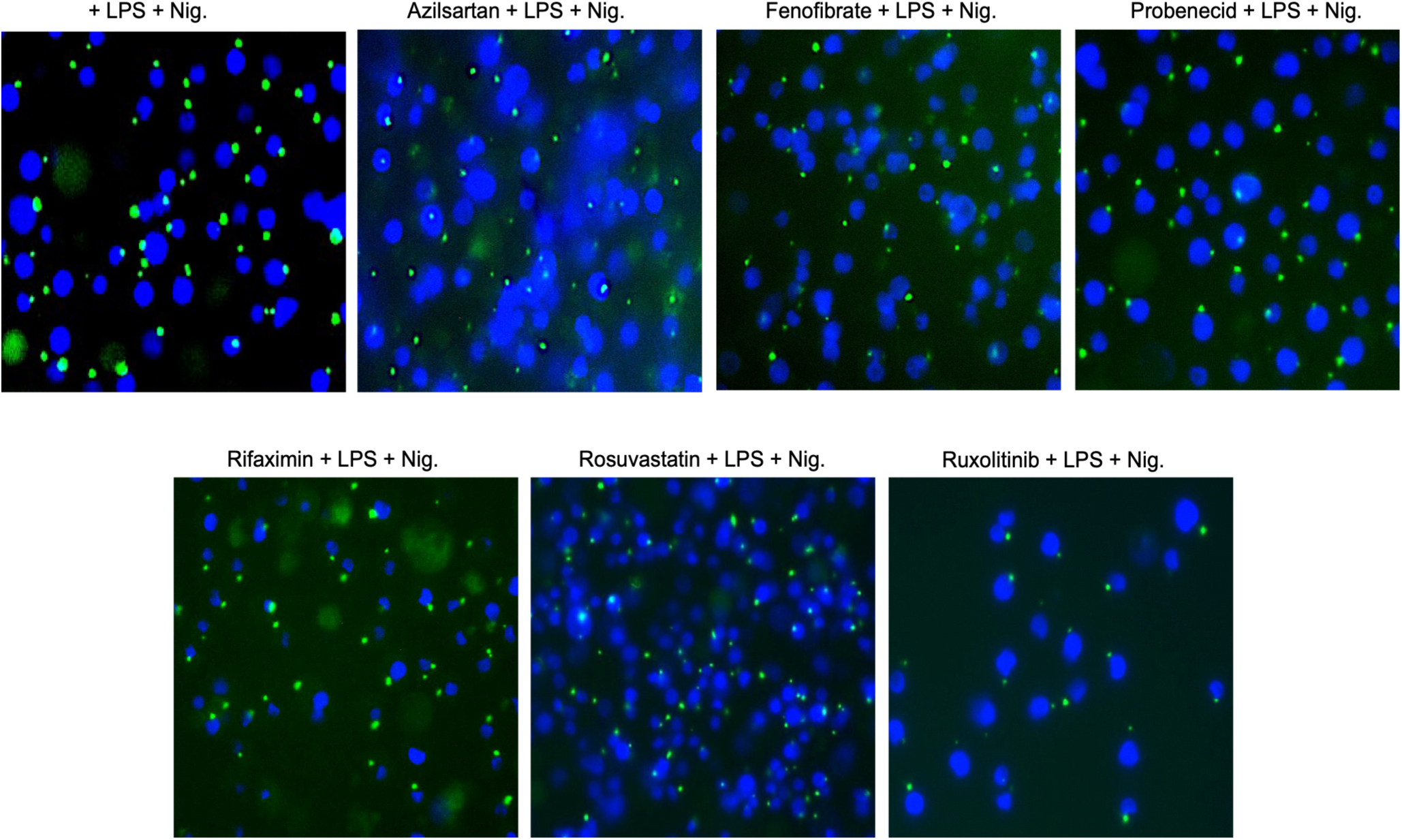
FDA-approved drugs showing 40-60% ASC-puncta reduction. ASC puncta formation in LPS + Nigericin treated THP-1-ASC-GFP cells with control vs. Azilsartan, Fenofibrate, Probenecid, Rifaximin, Rosuvastatin, and Ruxolitinib.

**Fig. S3:**
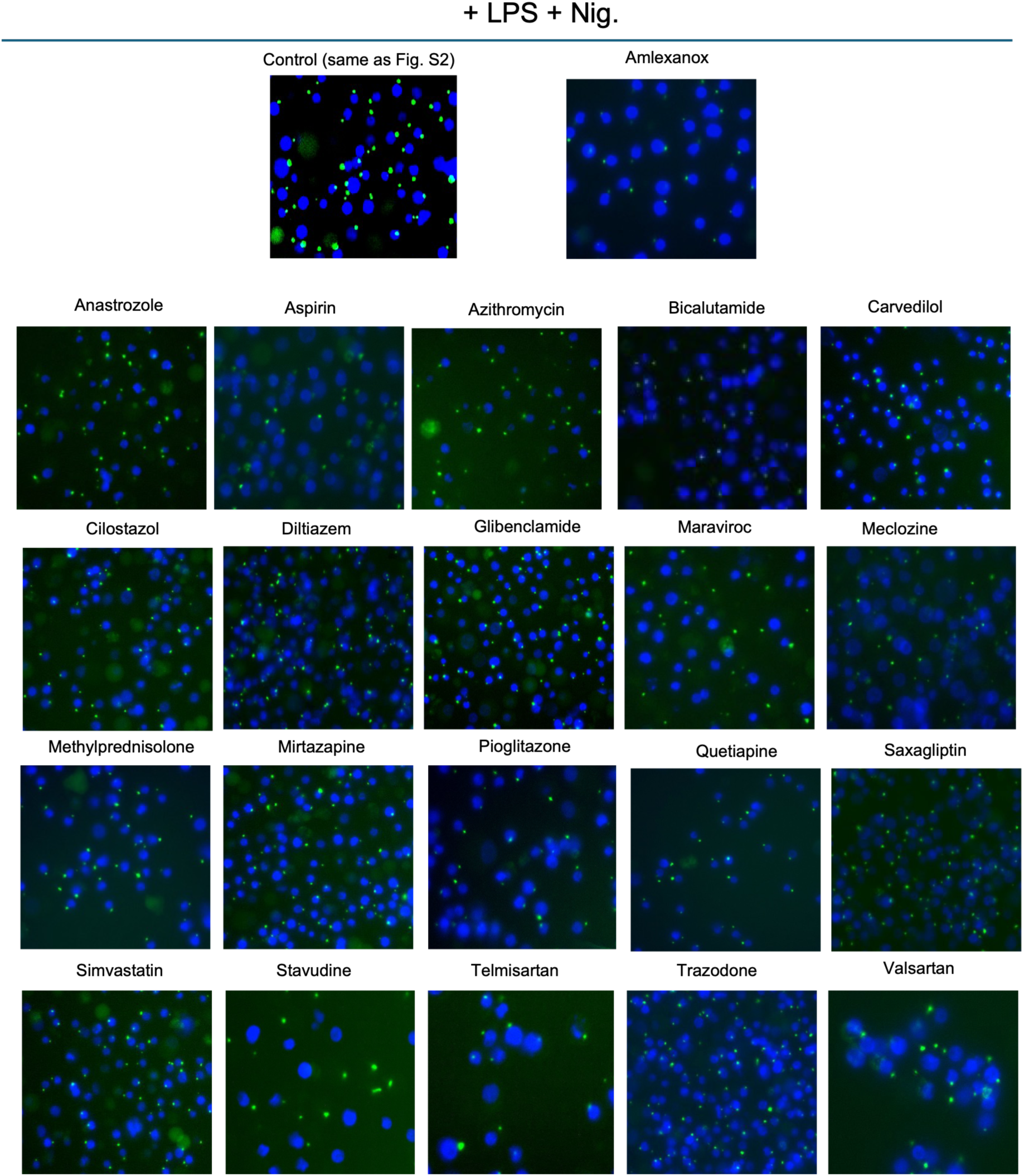
FDA-approved drugs showing 20-40% ASC-puncta reduction. ASC puncta formation in LPS + Nigericin treated THP-1-ASC-GFP cells with control vs. Amlexanox, Anastrozole, Aspirin, Azithromycin, Bicalutamide, Carvedilol, Cilostazol, Diltiazem, Glibenclamide, Maraviroc, Meclozine, Methylprednisolone, Mirtazapine, Pioglitazone, Quetiapine, Saxagliptin, Simvastatin, Stavudine, Telmisartan, Trazodone, and Valsartan.

**Fig. S4:**
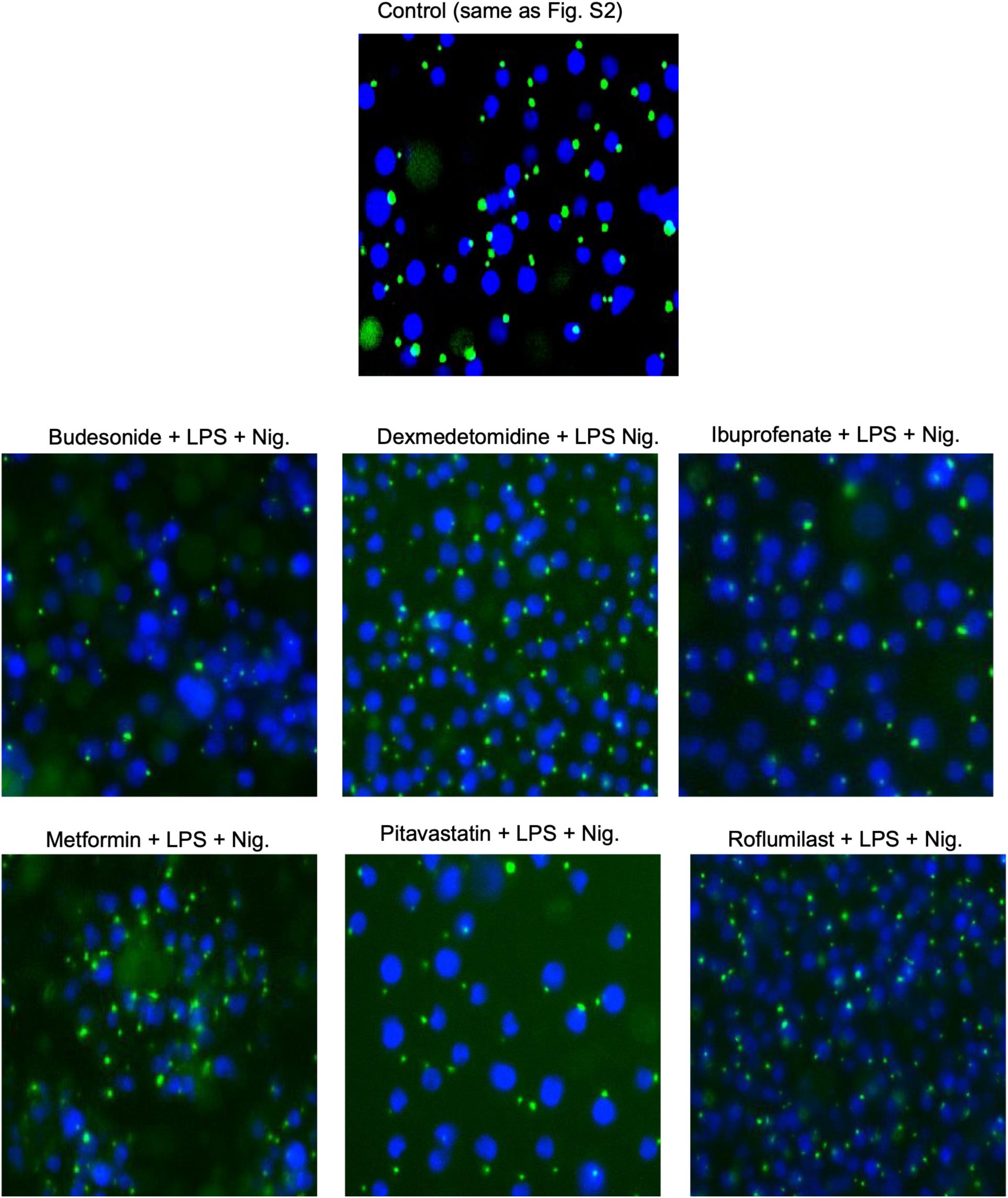
FDA-approved drugs showing 10-20% ASC-puncta reduction. ASC puncta formation in LPS + Nigericin treated THP-1-ASC-GFP cells with control vs. Budesonide, Dexmedetomidine, Ibuprofenate, Metformin, Pitavastatin, and Roflumilast.

**Fig. S5:**
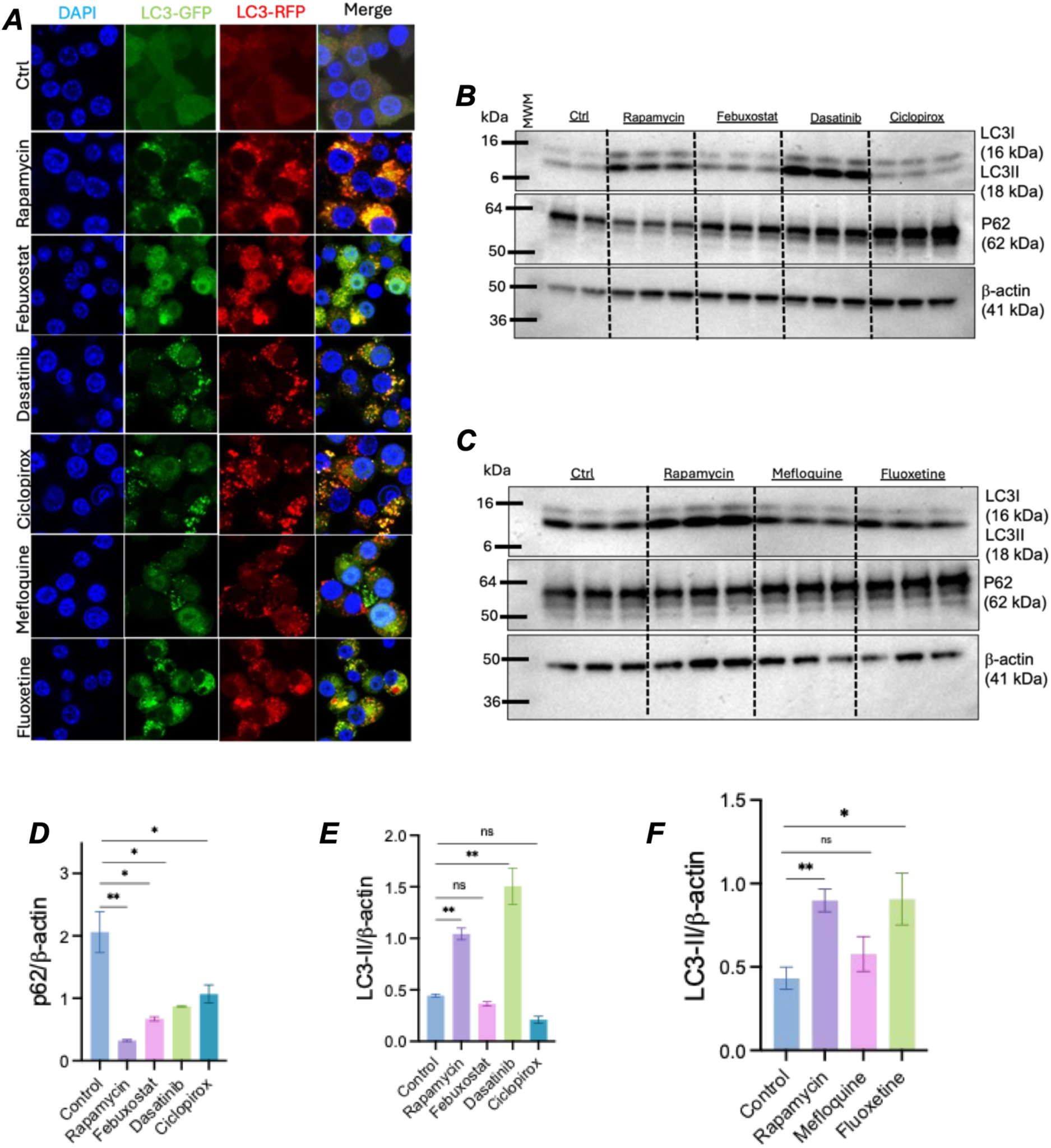
LPS priming blockers effect on autophagy. (***A***) Fluorescent microscopy of RAW-Difluo mLC3 cells treated with either saline of various drugs. (***B, C***) Western blot analysis of autophagy markers LC3-II and p62. (***D***) quantification of p62 vs β-actin. (***E, F***) quantification of LC3 vs β-actin.

